# Developing Cyclic Peptomers as Broad-Spectrum Gram negative Bacterial Type III Secretion System Inhibitors

**DOI:** 10.1101/2020.08.03.235622

**Authors:** Hanh N. Lam, Tannia Lau, Adam Lentz, Jessica Sherry, Alejandro Cabrera-Cortez, Karen Hug, Joanne Engel, R. Scott Lokey, Victoria Auerbuch

## Abstract

Antibiotic resistant bacteria are an emerging global health threat. New antimicrobials are urgently needed. The injectisome type III secretion system (T3SS), required by dozens of Gram-negative bacteria for virulence but largely absent from non-pathogenic bacteria, is an attractive antimicrobial target. We previously identified synthetic cyclic peptomers, inspired by the natural product phepropeptin D, that inhibit protein secretion through the *Yersinia* Ysc and *Pseudomonas aeruginosa* Psc T3SSs, but do not inhibit bacterial growth. Here we describe identification of an isomer, 4EpDN, that is two-fold more potent (IC_50_ 4 μM) than its parental compound. Furthermore, 4EpDN inhibited the *Yersinia* Ysa and the *Salmonella* SPI-1 T3SSs, suggesting that this cyclic peptomer has broad efficacy against evolutionarily distant injectisome T3SSs. Indeed, 4EpDN strongly inhibited intracellular growth of *Chlamydia trachomatis* in HeLa cells, which requires the T3SS. 4EpDN did not inhibit the unrelated Twin arginine translocation (Tat) system, nor did it impact T3SS gene transcription. Moreover, although the injectisome and flagellar T3SSs are evolutionarily and structurally related, the 4EpDN cyclic peptomer did not inhibit secretion of substrates through the *Salmonella* flagellar T3SS, indicating that cyclic peptomers broadly but specifically target the injestisome T3SS. 4EpDN reduced the number of T3SS basal bodies detected on the surface of *Y. enterocolitica*, as visualized using a fluorescent derivative of YscD, an inner membrane ring with low homology to flagellar protein FliG. Collectively, these data suggest that cyclic peptomers specifically inhibit the injectisome T3SS from a variety of Gram-negative bacteria, possibly by preventing complete T3SS assembly.

**IMPORTANCE:** Traditional antibiotics target both pathogenic and commensal bacteria, resulting in a disruption of the microbiota, which in turn is tied to a number of acute and chronic diseases. The bacterial type III secretion system (T3SS) is an appendage used by many bacterial pathogens to establish infection, but is largely absent from commensal members of the microbiota. In this study, we identify a new derivative of the cyclic peptomer class of T3SS inhibitors. These compounds inhibit the T3SS of the nosocomial ESKAPE pathogen *Pseudomonas aeruginosa* and enteropathogenic *Yersinia* and *Salmonella*. The impact of cyclic peptomers is specific to the T3SS, as other bacterial secretory systems are unaffected. Importantly, cyclic peptomers completely block replication of *Chlamydia trachomatis*, the causative agent of genital, eye, and lung infections, in human cells, a process that requires the T3SS. Therefore, cyclic peptomers represent promising virulence blockers that can specifically disarm a broad spectrum of Gram-negative pathogens.

## INTRODUCTION

Antibiotic resistance is of great concern to global public health. Bacterial pathogens have evolved numerous mechanisms to survive treatment with clinically-available antibiotics (1). Alternative therapies against multidrug-resistant strains of so-called ESKAPE pathogens (*<u>E</u>nterococcus faecium*, *<u>S</u>taphylococcus aureus*, *<u>K</u>lebsiella pneumoniae*, *<u>A</u>cinetobacter baumannii*, *<u>P</u>seudomonas aeruginosa*, and *<u>E</u>nterobacter* species) are urgently needed. Various strategies have been explored to avoid the antimicrobial apocalypse (2). One promising approach is to inhibit bacterial virulence mechanisms to disarm pathogens without affecting non-pathogenic members of the microbiota or environmental bacteria (3, 4). This approach has the potential to not only control infection but to do so in a way that preserves the integrity of the microbiome, which is beneficial for human health and is often the source of antibiotic resistance genes (5, 6).

The type III secretion system (T3SS), a needle-like injectisome apparatus, is required for virulence in many Gram-negative pathogens including *Salmonella*, *Yersinia*, *Chlamydia* and the ESKAPE pathogen, *P. aeruginosa*. The T3SS is largely absent from commensal bacteria, making it a good target for virulence blocker antimicrobials. Phylogenetic analysis suggests that T3SSs evolved from the flagellar system (7, 8). Indeed, the flagellar basal body is a secretion system, referred to as the flagellar T3SS, that secretes flagellin and other structural components into the extracellular space in order to build the flagellar filament to power motility. The flagellar and injectisome T3SS share a number of conserved basal body and export apparatus components (9). However, the injectisome T3SS does not mediate motility, but instead delivers effector proteins into target host cells.

A number of small molecules, antibodies, and vaccines have been studied for T3SS targeted therapies (10). Despite showing promising effects on the T3SS *in vitro* and in animal models, only one antibody-based therapy has entered clinical trials. A bispecific antibody, MEDI3902, against the *P. aeruginosa* T3SS needle tip protein PcrV and the Psl exopolysaccharide is effective against both acute and chronic infection models and is in phase II clinical trials for prevention of ventilator-associated pneumonia (11, 12).

As narrow-spectrum antimicrobials require more precise diagnostics, broad-spectrum T3SS inhibitors would be more valuable clinically than those only able to target one bacterial species. In addition, most mammalian pathogens that utilize a T3SS only require their T3SS during growth within, but not outside, the host animal. However, *Chlamydiae,* which cause lung, genital, and eye infections, are obligate intracellular bacteria and their T3SS is strictly required for their growth (13). Interestingly, the *Chlamydia* T3SS belongs to its own T3SS family (7, 8). Here we identify a derivative of a synthetic cyclic peptomer family of T3SS inhibitors (14) that can inhibit the T3SS machinery of three evolutionarily distinct T3SS families used by five different bacterial species to cause human disease, including *Chlamydia trachomatis*.

## RESULTS

### Structure-activity relationship study of cyclic peptomers

Previously we identified a group of cyclic peptomers that inhibited secretion of substrates from *Y. pseudotuberculosis* and *P. aeruginosa* T3SSs, but did not inhibit bacterial growth, motility, or HeLa cell metabolism (14). The results suggested a potential for development of these cyclic peptomers as pathogen-specific virulence blockers. Based on dose response curves and concentration of half maximal inhibition (IC_50_) of the *P. aeruginosa* T3SS, 1EpDN (previously named as EpD1,2N) was chosen for structure-activity relationship (SAR) analysis. Compounds used in SAR analysis are listed in Table 1.

**Table 1:**
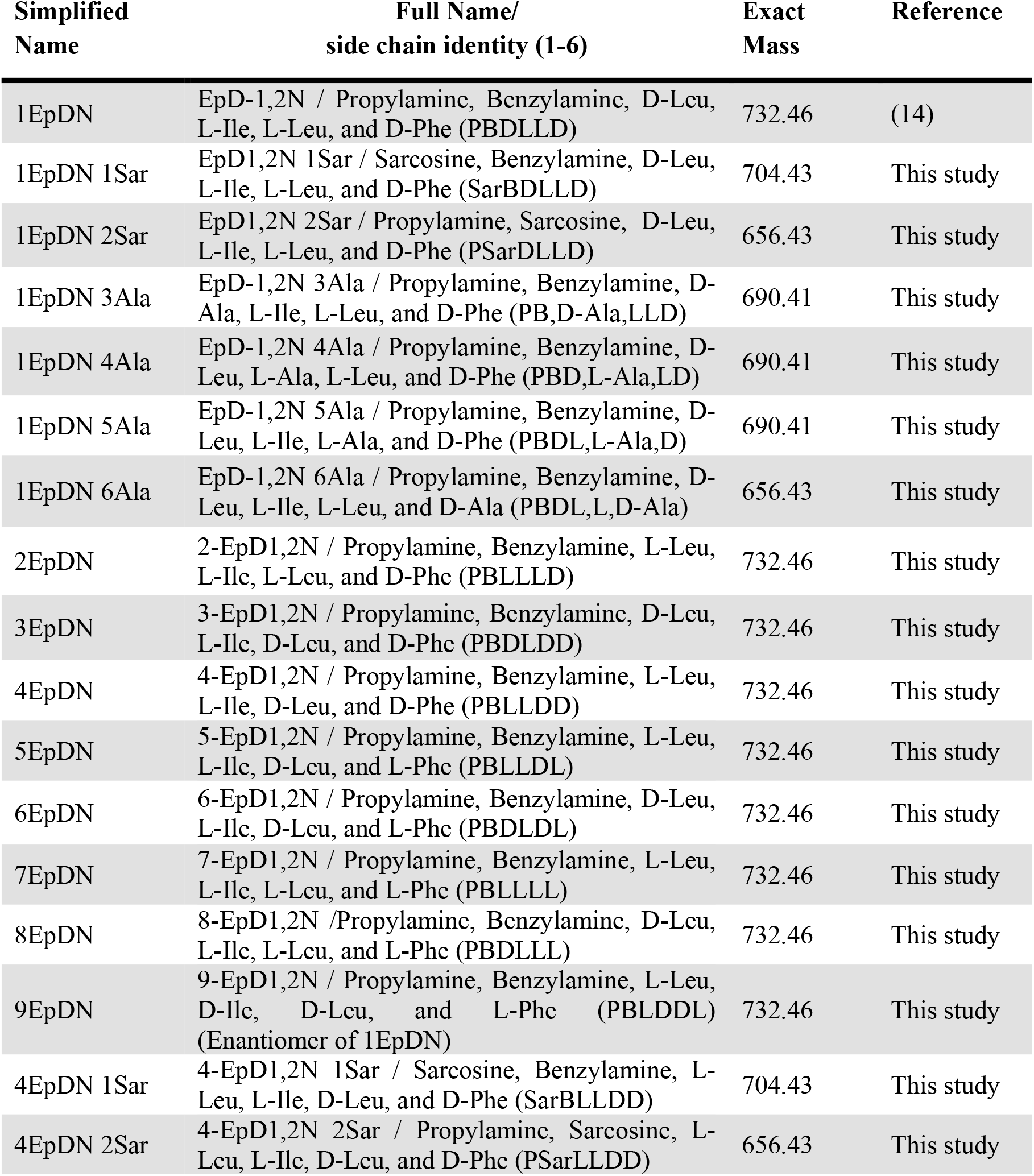
Compounds synthesized and used in this study.

We first assessed the effect of alanine replacement at each of the six positions of the parent scaffold, 1EpDN. Note that because peptoids have side chains appended to a nitrogen atom rather than carbon as in amino acids, positions 1 and 2 were synthesized with N-methylglycine, also known as sarcosine (Sar), as the peptoid equivalent of alanine (Ala). Ala or Sar replacement at any of the six positions resulted in significant loss of activity, suggesting that all side chains contribute to the activity (Fig. S1). Next, we carried out a stereochemistry scan, in which different combinations of L- and D-amino acids at positions 3 to 6 were generated. The parent compound, 1EpDN, has <u>**P**</u>ropylamine, and <u>**B**</u>enzylamine at positions 1 and 2, and <u>**D**</u>-Leu, <u>**L**</u>-Ile, <u>**L**</u>-Leu, and <u>**D**</u>-Phe at positions 3-6. For the stereochemistry scan, we will refer to 1EpDN as PBDLLD.

While most stereoisomers had the same or reduced T3SS inhibitory activity, 4EpDN (PBLLDD) showed improved activity, with an IC_50_ of ~4μM compared to the parent compound IC_50_ of ~8μM (Fig. 1A-B). Replacement of position 1 (4EpDN 1Sar) or position 2 (4EpDN 2Sar) with Sar significantly reduced activity of 4EpDN (Fig. 2A-B). 4EpDN and 4EpDN 2Sar were used as an active compound and a negative control, respectively, in most follow-up experiments.

**Figure 1:**
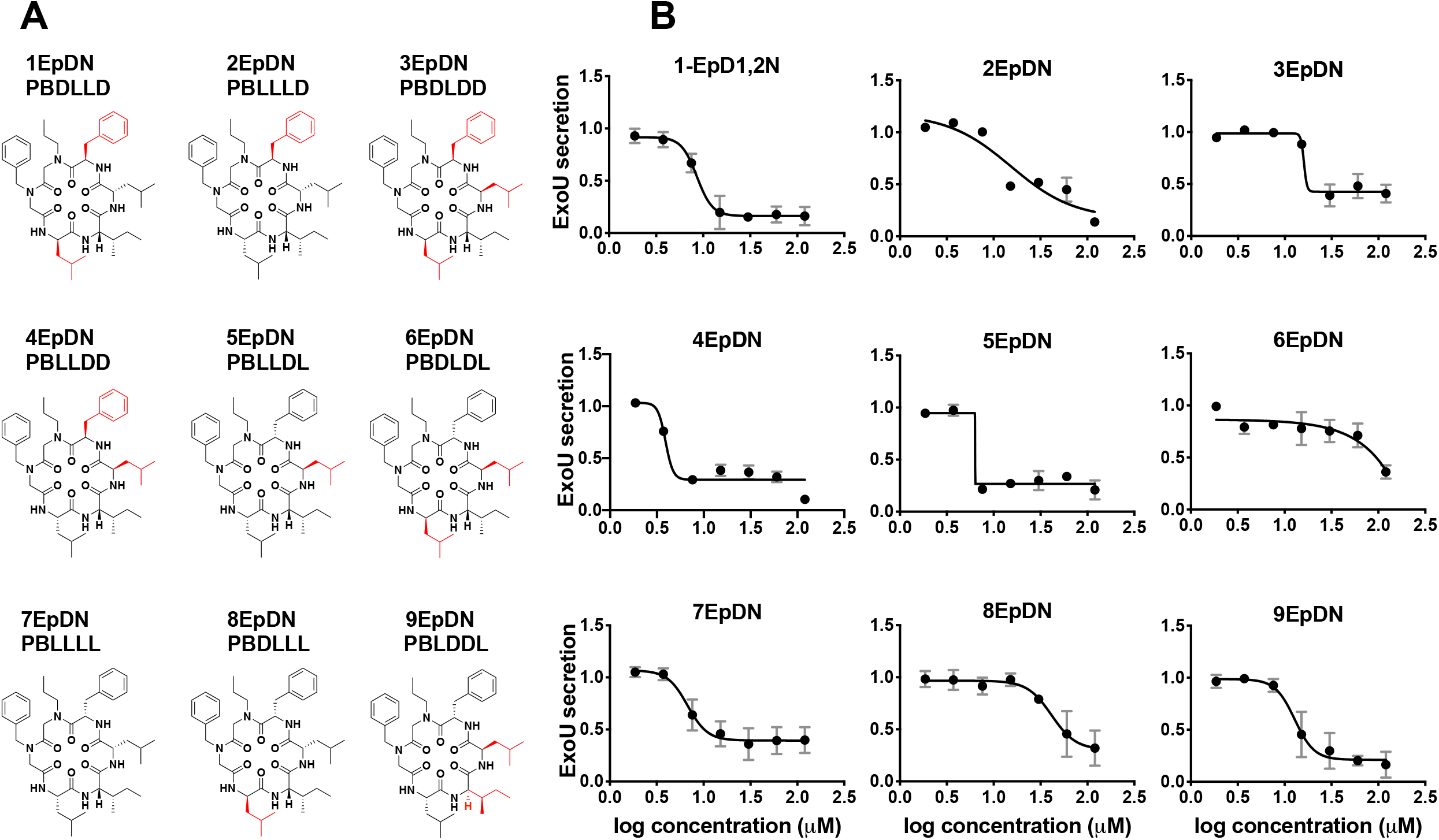
Stereochemistry scan of cyclic peptomers results in a more potent derivative, 4EpDN. **(A)** Structures of 1EpDN stereoisomers. Isomers were generated from different combination of four side chains at position 3 to 6. Numbers preceding compounds were used to distinguish the different isomers and the conformation of the four side chains. D-amino acid side chain is shown in red. **(B)** WT *P. aeruginosa* PA103 was grown under T3SS-inducing conditions with increasing concentrations of cyclic peptomer isomers. Secretion of T3SS cargo into the culture supernatant was assessed by precipitating secreted proteins and visualizing them with Coomassie blue. ExoU band intensities were quantified and normalized to that of the DMSO control. The results are from at least two independent experiments. Nonlinear curve fitting is shown to depict the trend of inhibition.

**Figure 2:**
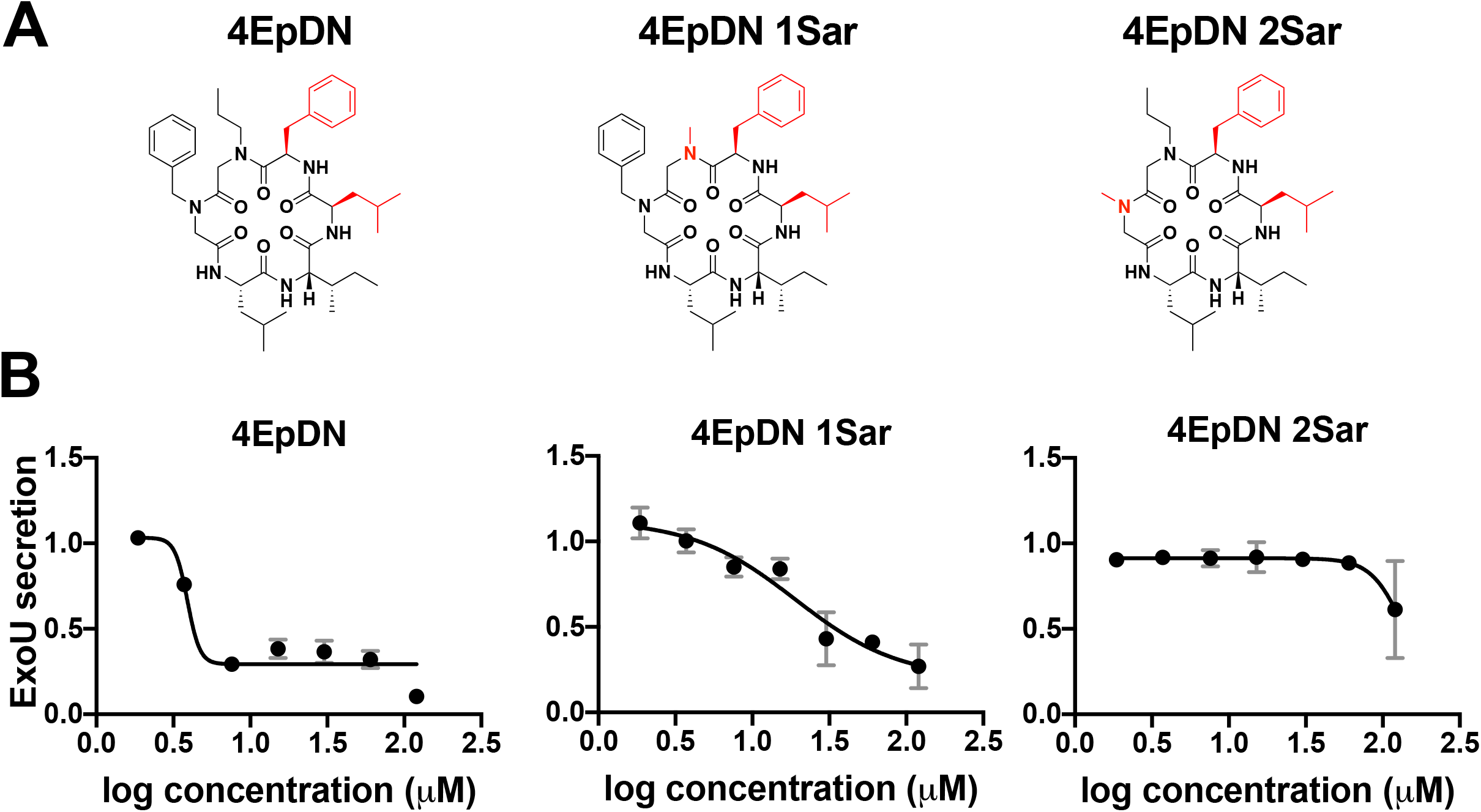
Sarcosine replacement of 4EpDN at position 1 or 2 eliminates activity. **(A)** Structures of 4EpDN and its derivatives, 4EpDN 1Sar and 4EpDN 2Sar. D-amino acid side chain is shown in red. **(B)** WT *P. aeruginosa* PA103 was grown under T3SS-inducing conditions with increasing concentrations of compounds. Secretion of T3SS cargo into the culture supernatant was assessed on SDS-PAGE gel. ExoU band intensities were visualized with Coomassie blue, quantified and normalized to that of the DMSO control. The results are from at least two independent experiments.

### Cyclic peptomers inhibit secretion of T3SS substrates from the Inv-Mxi-Spa T3SS family, but does not inhibit secretion through the flagellar T3SS

Based on phylogenetic analysis of core T3SS proteins, T3SSs were classified into seven families (7, 8). However, T3SSs have many highly conserved structural components (9). T3SS genes are typically encoded on virulence plasmids or pathogenicity islands, indicative of horizontal gene transfer (15); therefore, phylogeny of T3SSs does not follow organismal phylogeny. We previously showed that cyclic peptomers inhibited the Ysc T3SS family found in *P. aeruginosa* and *Yersinia* (Fig. 1-2) (14). In order to test whether cyclic peptomers are active against other T3SS families, we evaluated the effect of cyclic peptomers on the Inv-Mxi-Spa T3SS in *Y. enterocolitica* and *Salmonella enterica* serovar Typhimurium.

The *Y. enterocolitica* Ysa system, a chromosomally encoded T3SS, is distinct from the *Yersinia* Ysc T3SS and contributes to *Y. enterocolitica* colonization of the terminal ileum and gastrointestinal system associated tissues (16, 17). A *Y. enterocolitica* mutant that lacks expression of the Ysa T3SS (Δ*ysaT*) was used as a negative control, while a mutant lacking the Ysc T3SS (Δ*yscL*) (18) was used to evaluate the effect of compounds specifically on the Ysa system. Secretion of the Ysa effector protein YspF was quantified. 4EpDN inhibited secretion of YspF in a dose dependent manner, while 4EpDN 2Sar did not affect its secretion (Fig. 3). Together, these results suggest that cyclic peptomers are active against both the Ysc and Ysa T3SSs in *Yersinia*.

**Figure 3:**
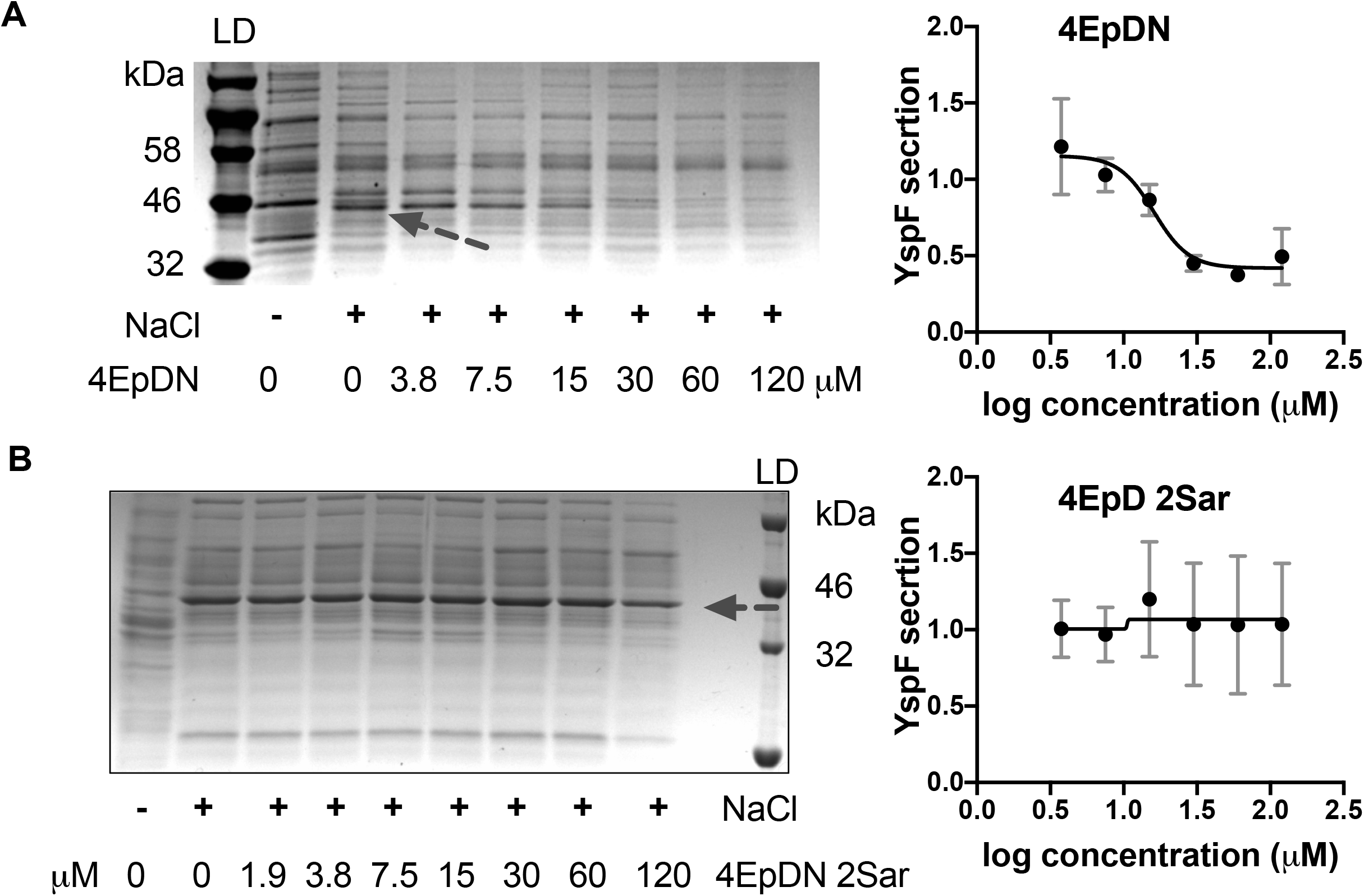
Effect of cyclic peptomers on secretion of *Yersinia* Ysa T3SS substrates. Y *. enterocolitica* serotype O:8 was grown under T3SS-inducing conditions with increasing concentrations of cyclic peptomer isomers, 4EpDN **(A)** and 4EpDN 2Sar **(B)**. Secretion of T3SS cargo into the culture supernatant was assessed by precipitating secreted proteins and visualizing them with Coomassie blue. YspF band intensities were quantified and normalized to that of the DMSO control. Representative gel images and quantification of YspF are shown. The results are from two independent experiments.

In order to evaluate whether the cyclic peptomers are active against T3SSs distinct from the Ysc T3SS outside the *Yersinia* genus, we tested cyclic peptomer efficacy in *Salmonella*. *Salmonella* employs two T3SSs during infection, with the SPI-1 T3SS belonging to the Inv-Mxi-Spa T3SS family (7, 8). Inhibition of SPI-1 T3SS effector protein SipC and SipA (19–21) secretion by 4EpDN was observed at ~1 μM and ~1.4 μM, respectively, while 4EpDN 2Sar showed inhibition of SipC and SipA secretion only at concentrations greater than 30 μM (Fig. 4). In order to test whether inhibition of secretion by the cyclic peptomers in *Salmonella* could be affected by aggregation of the compounds, we also evaluated secretion in the presence of detergent. The presence of Tween-20 (0.003%, Fig. S2) did not reduce activity of 4EpDN (Fig. 4), suggesting that aggregation of cyclic peptomers does not affect its activity.

**Figure 4:**
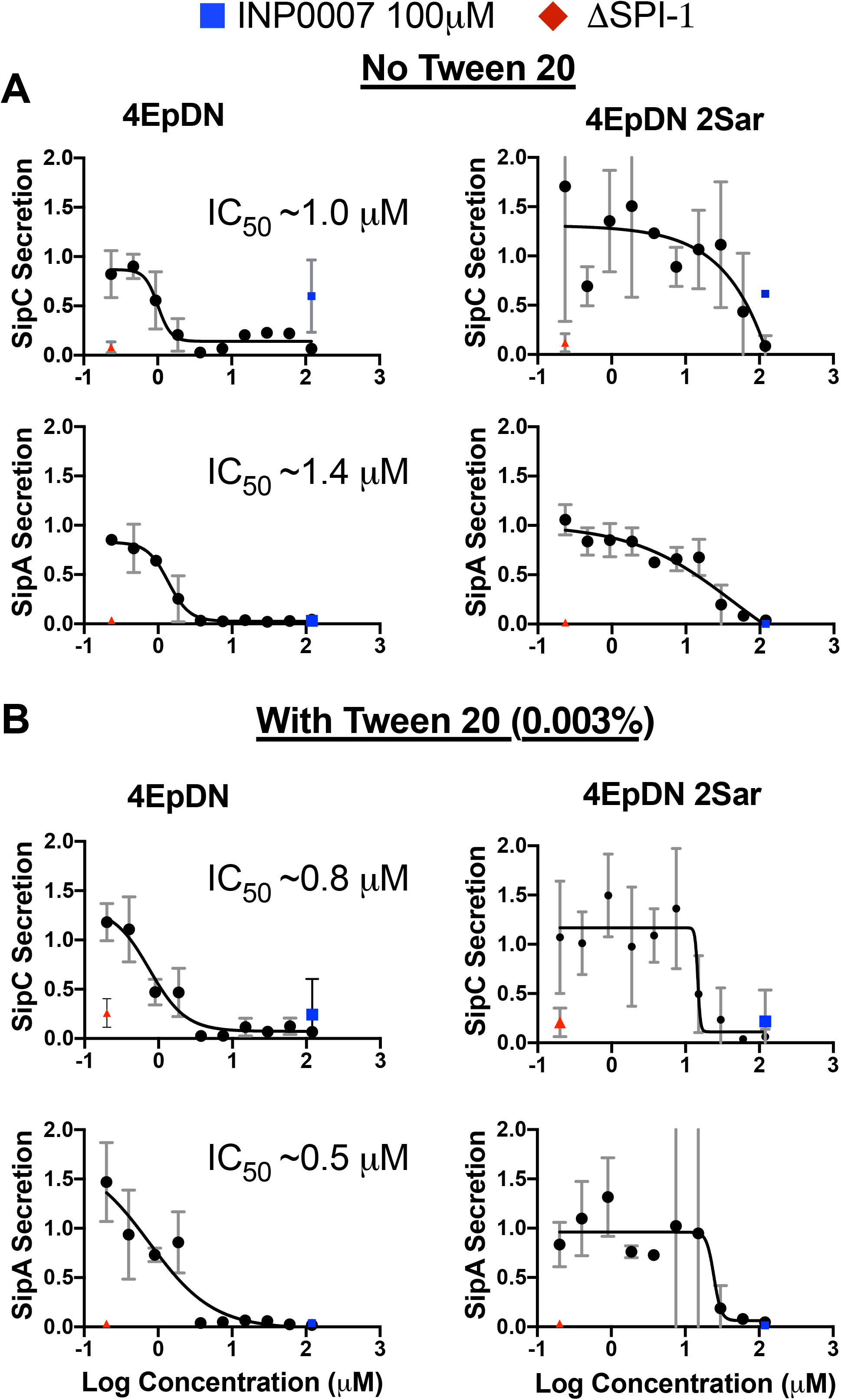
Cyclic peptomers inhibit the *Salmonella* SPI-1 T3SS. *Salmonella enterica* Typhimurium was grown with increasing concentrations of cyclic peptomer isomers. Secretion of SPI-1 T3SS cargo into the culture supernatant was assessed by precipitating secreted proteins and visualizing them with Coomassie blue. SipA and SipC band intensities were quantified and normalized to that of the DMSO control. The experiments were carried out without the detergent Tween 20 **(A)**, or with Tween 20 **(B).** A ΔSPI-1 *Salmonella* mutant and INP0007, a known SPI-1 inhibitor (60), were used as controls. The results are from at least two independent experiments.

As 4EpDN inhibited both the Ysc and Inv-Mxi-Spa T3SS families, we tested whether this cyclic peptomer could inhibit the flagellar T3SS, which is the most distantly related T3SS based on previous phylogenetic analysis (7). Conveniently, *Salmonella* expresses the SPI-1 and its flagellar system under the same conditions *in vitro* (rich media). This allowed us to investigate effects of cyclic peptomers on both the SPI-1 T3SS and flagellar systems under the same culture conditions. Because of the conservation between the injectisome and flagellar T3SSs, flagellar substrates can be secreted through both systems. Therefore, secretion of flagellar substrates (FliC and FliD) was quantified in both WT and ΔSPI-1 strains to distinguish secretion through both the SPI-1 T3SS and flagellar system (WT strain) or only through the flagellar system (ΔSPI-1 strain). 4EpDN inhibited FliC and FliD secretion in WT *Salmonella* at concentrations of ≥60 μM and ≥3.75 μM, respectively (Fig. S3), consistent with the ability of the SPI-1 T3SS being able to secrete flagellar substrates. However, 4EpDN only inhibited FliC and FliD secretion at high concentrations (≥ 60μM) in the ΔSPI-1 mutant, with unfavorable dose response curves compared to WT *Salmonella*. This suggests that the inhibitory effect of 4EpDN on FliD secretion in the WT strain was mainly through inhibition of its secretion through the SPI-1 T3SS. 4EpDN 2Sar had no significant effect on FliC secretion or FliD secretion. These data suggest that the cyclic peptomer 4EpDN does not significantly inhibit substrate secretion through the flagellar T3SS in *Salmonella* but strongly inhibits the SPI-1 T3SS under the same conditions.

### Cyclic peptomers affect the T3SS basal body

The T3SS basal body must be assembled prior to T3SS substrate secretion (22–24). In *Yersinia*, the T3SS basal body component YscD is an inner ring protein that is conserved among injectisome T3SSs, but has low sequence homology with the flagellar ortholog FliG (9). Absence of YscD at the inner membrane prevents assembly of other T3SS machinery (YscL, YscK, YscQ) (23, 25), and secretion of T3SS substrates (26). We used a *Y. enterocolitica* strain expressing a YscD allele translationally fused with EGFP to visualize the effect of compounds on YscD assembly (23). 4EpDN caused reduction in the number of YscD puncta, although not to the levels seen under non-T3SS inducing conditions (high Ca^2+^), while the inactive isomer 4EpDN 2Sar had no effect (Fig. 5). These data suggest that cyclic peptomers affect the assembly or stability of the T3SS basal body, ultimately dampening secretion of effector proteins.

**Figure 5:**
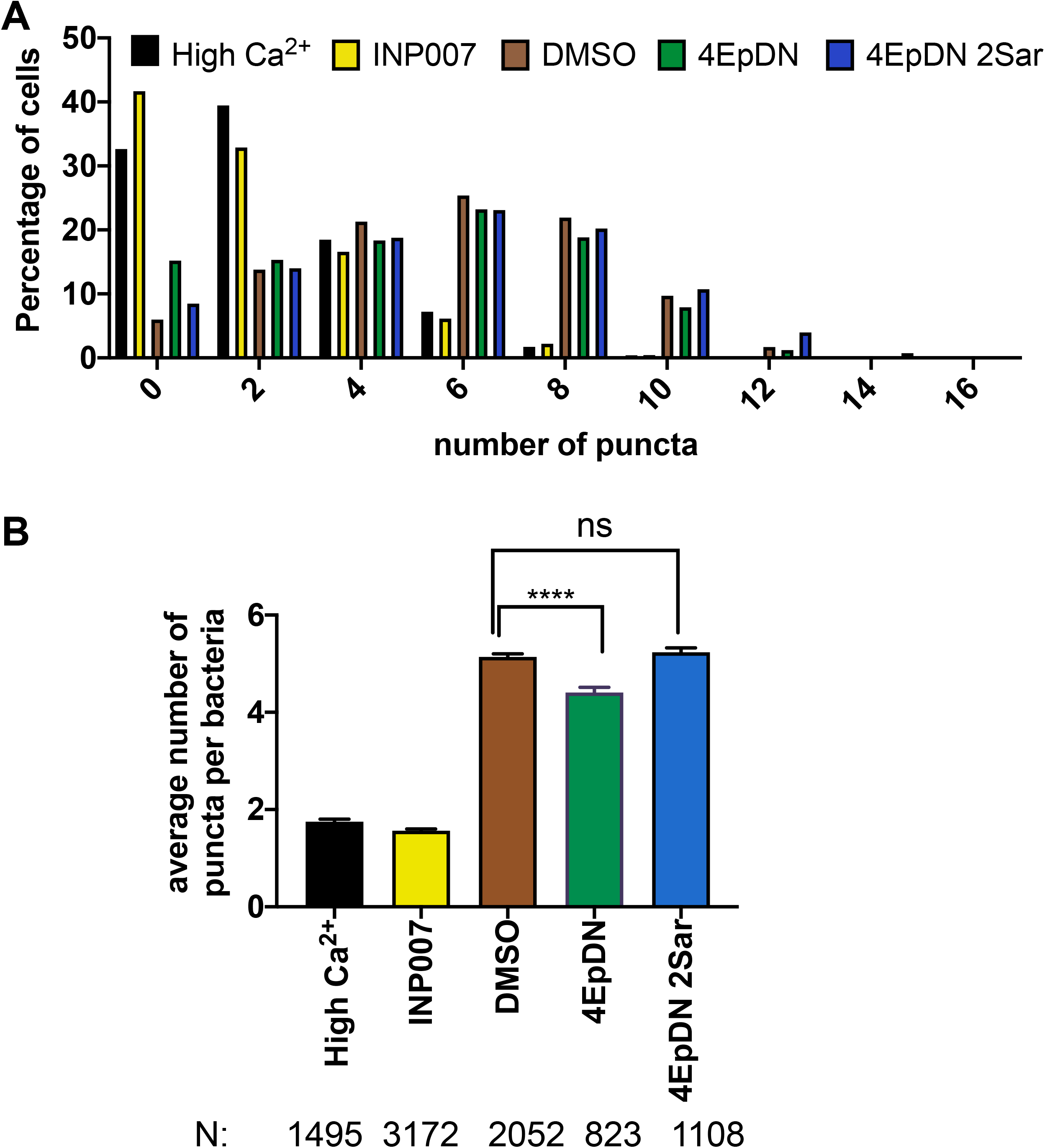
The cyclic peptomer 4EpDN disrupts localization of the *Yersinia* T3SS basal body component YscD. *Y. enterocolitica* expressing YscD-EGFP was grown under T3SS inducing condition (low Ca^2+^) in the presence of 9 μM cyclic peptomers, 50 μM INP007, or DMSO. High Ca^2+^ media was used as a non-secreting control. **(A)** Histogram showing the frequency of YscD puncta/cell. **(B)** Average number of puncta/cell after treatment ± standard error of the mean. Data represents three independent experiments. One-way ANOVA with Dunnett’s multiple-comparison test was used. ****, P < 0.0001; ns: not significant.

### Cyclic peptomers do not inhibit T3SS gene expression

As cyclic peptomers inhibit substrate secretion and basal body assembly on the bacterial membrane, we evaluated the effect of cyclic peptomers on T3SS gene expression, as this step occurs prior to T3SS assembly. Interestingly, 1EpDN did not inhibit transcription of *exoT* while phenoxyacetamide MBX1641 did (Fig. 6A). Expression of the *dsbA* gene, whose product was shown to be required for expression of the T3SS (27), was unchanged following either cyclic peptomer or the phenoxyacetamide treatment (Fig. 6A). Similarly, 4EpDN did not affect expression of the effectors *exoS* and *exoT* in a strain of *P. aeruginosa* lacking all known efflux pumps (28) (Fig. 6B). In contrast, MBX1641 strongly reduced expression of both *exoS* and *exoT* in this strain. These data suggest that cyclic peptomers do not block T3SS activity by blocking T3SS expression.

**Figure 6:**
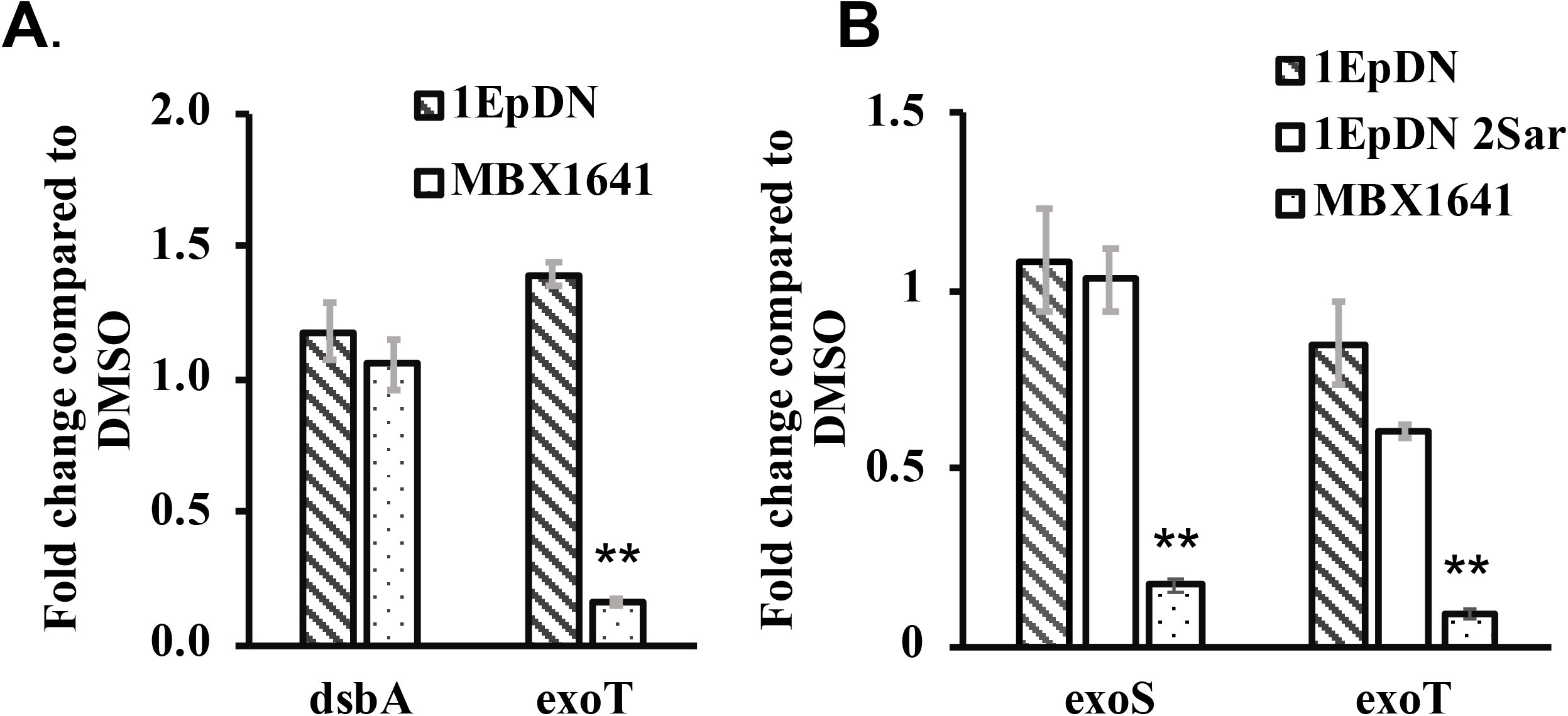
Cyclic peptomers do not affect transcription of T3SS genes in *P. aeruginosa*. *P. aeruginosa* PA103 **(A)** or PA01 **(B)** was grown in low calcium media in the presence of 60μM cyclic peptomers or DMSO. Samples were collected 3 hrs after induction for qPCR analysis. The phenoxyacetamide MBX1641 (53), a known T3SS inhibitor predicted to inhibit type III secretion by binding to the T3SS needle subunit (28), was used as a control. Data were from three replicates, analyzed by one-way ANOVA with Dunnett’s multiple-comparison test. **, P < 0.01.

In *P. aeruginosa*, secretion and transcription are coupled through a negative regulator ExsE (29). Secretion of ExsE allows transcription of T3SS genes by the *Pseudomonas* regulator ExsA (29). We evaluated the impact of cyclic peptomers on secretion of ExsE and the effector ExoS by Western Blot. Both 1EpDN and 4EpDN significantly reduced the amount of secreted ExoS, with a concomitant accumulation of ExoS in the bacterial cytosol (Fig. 7). In contrast, we did not observe any decrease in secreted ExsE and could not detect any cytosolic ExsE (data not shown). These data suggest that cyclic peptomers block effector protein secretion but not secretion of T3SS regulators.

**Figure 7:**
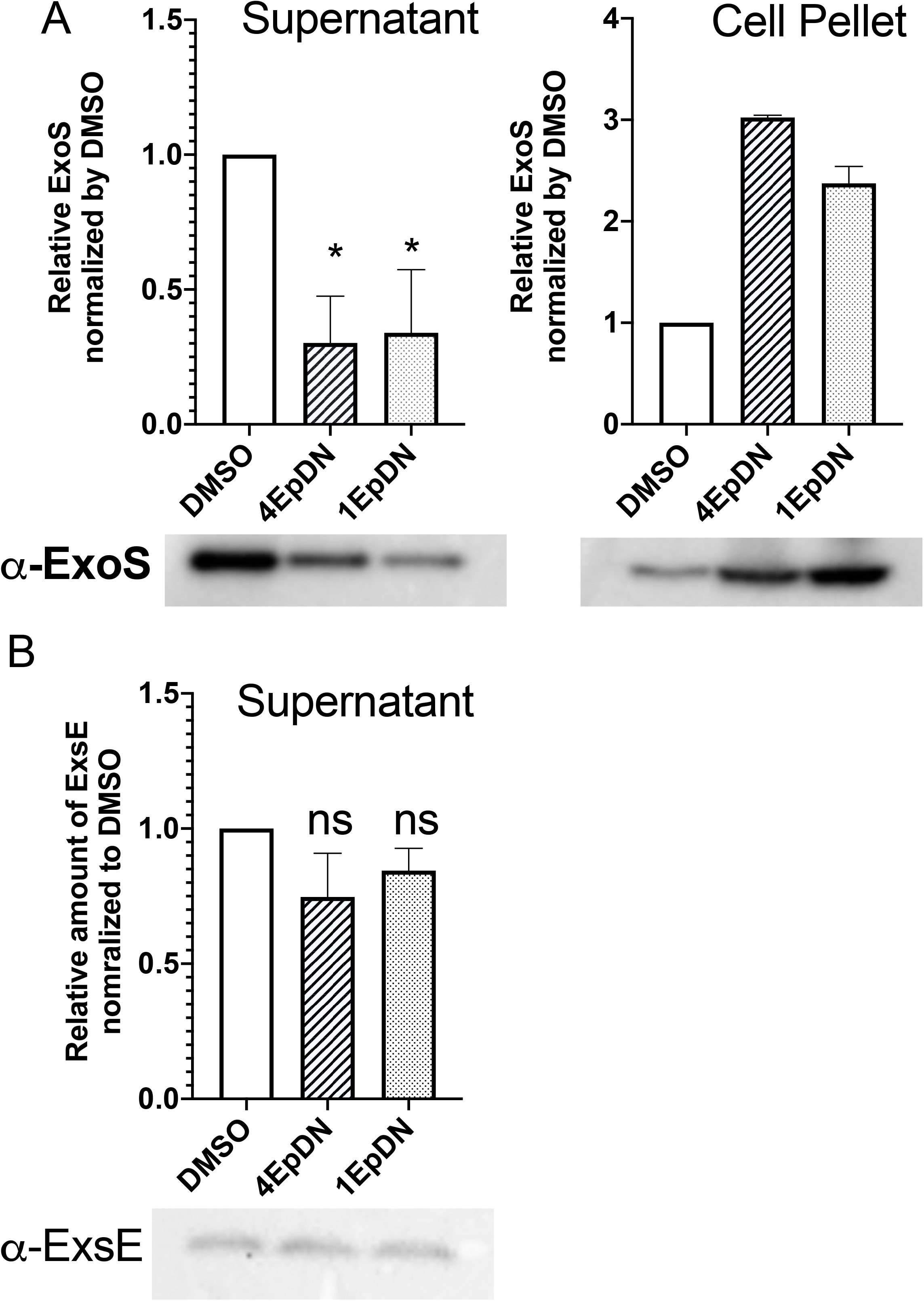
The cyclic peptomer 4EpDN inhibits secretion of the effector protein ExoS, but not the regulator ExsE. PAO1 carrying ExoS-Bla was grown in T3S inducing condition on the presence of 60μM cyclic peptomers or DMSO. **A**. Secretion of ExoS into the culture supernatant and synthesis of ExoS in the cell pellets were assessed by Western Blot using a β-lactamase antibody. **B**. In the same samples, Western blot was carried out for ExsE. ExsE in the supernatant was observed and quantified while ExsE in the cell pellets was undetectable. Data were from at least two independent experiments. One-way ANOVA with Dunnett’s multiple-comparison test was used. *, P < 0.05; ns: not significant, compared to DMSO.

### Cyclic peptomers do not inhibit secretion through the Twin-arginine translocation (Tat) system

In order to determine if cyclic peptomers inhibit the activity of secretion systems completely unrelated to the T3SS, we sought to assess the impact of cyclic peptomers on the Twin arginine translocation (Tat) system in *Y. pseudotuberculosis*. The Tat system translocates fully folded substrates across the inner membrane, while the T3SS translocates partially unfolded substrates across the inner, outer, and target host cell membranes (30). To monitor Tat secretion system activity, a reporter strain expressing an IPTG-inducible β-lactamase TEM-1 domain fused to the signal peptide of the SufI Tat substrate (31) was constructed. Following IPTG induction, β-lactamase confers resistance to the β-lactam peptidoglycan-targeting antibiotic penicillin G when the SufI-β-lactamase reporter has successfully translocated into the periplasm (Fig. 8A). The presence of known Tat inhibitors, Bay 11-7082 or N-phenylmaleimide (32), strongly reduced growth of bacteria after four and six hours, while growth of bacterial cultures treated with cyclic peptomers were similar to the DMSO control (Fig. 8B,C). These results suggested that 4EpDN does not inhibit the Tat secretion system.

**Figure 8:**
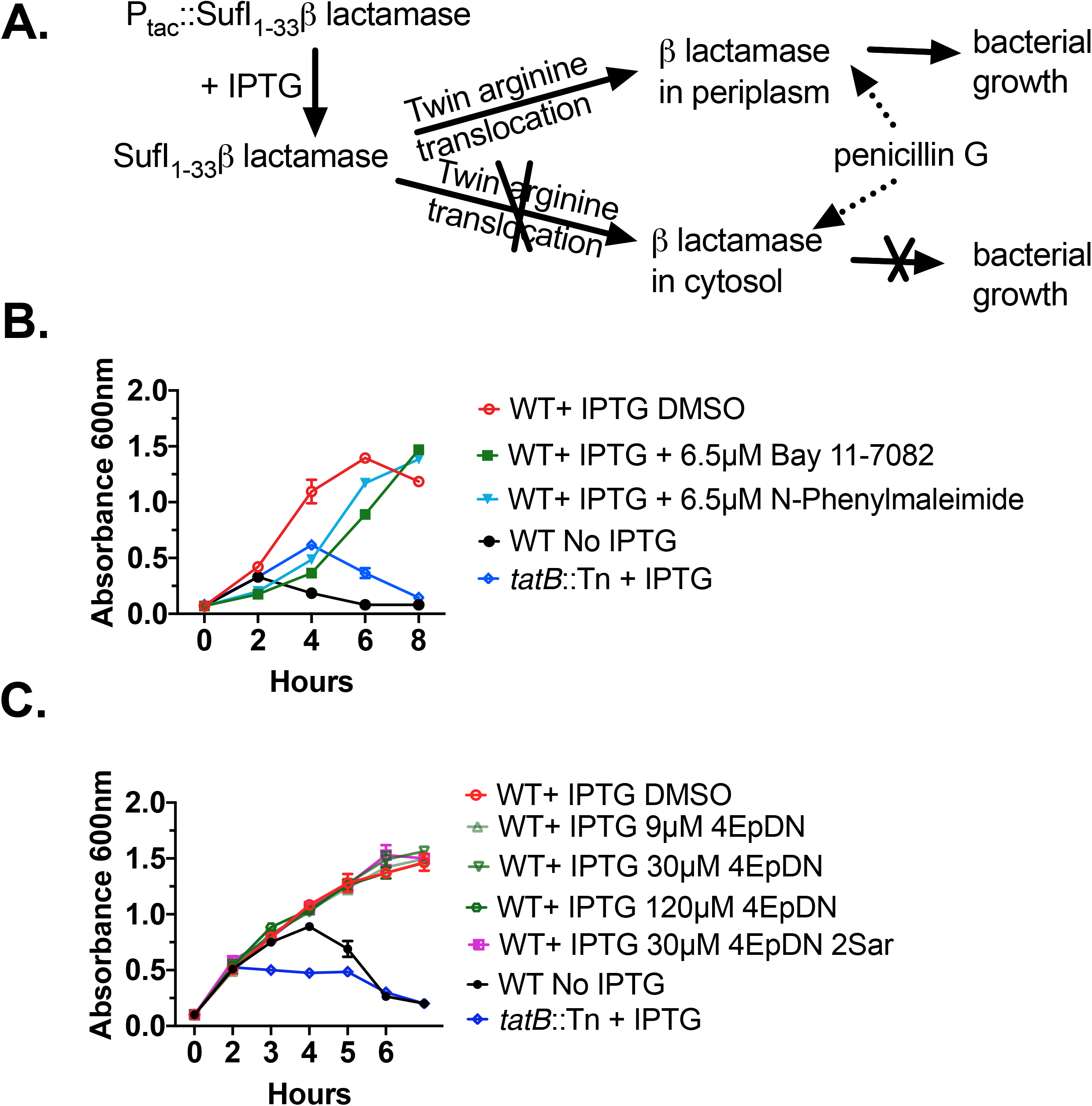
Cyclic peptomers do not affect the twin arginine translocation (Tat) system. **(A)** *Y. pseudotuberculosis* expressing a SufI-β-lactamase Tat secretion reporter incubated in the presence of penicillin G will only grow if the Tat secretion system remains functional. **(B)** *Y. pseudotuberculosis* SufI-β-lactamase reporters were treated with the Tat inhibitors Bay 11-7082, N-Phenyl maleimide, or DMSO, and culture optical density was measured. WT refers to bacteria expressing a functional Tat secretion system. A mutant strain with a transposon insertion in the *tatB* gene serves as a control. **(C)** The same assay as in (B) was repeated in the presence of cyclic peptomers or DMSO. The result was from two independent replicates.

### Cyclic peptomers block *Chlamydia* infection

Our data point to the cyclic peptomer 4EpDN being able to inhibit multiple T3SS injectisome families but not other secretion systems, indicating a broad yet specific activity. In order to evaluate whether this cyclic peptomer can disarm virulence, we chose to examine the effect of this compound on the *Chlamydia* infection, as this pathogen requires the T3SS for infection and growth within human cells. The *Chlamydial* life cycle involves two major bacterial forms: the extracellular infectious Elementary Bodies (EBs), and the intracellular replicative Reticulate Bodies (RBs). Upon entry, EBs discharge preloaded T3SS effectors and are taken up into a membrane-bound compartment (the inclusion) where they differentiate into RBs, secrete additional T3SS effectors and replicate, and then re-differentiate into EBs. Initial stages of infection were assessed by quantifying the number of inclusions/cell at 24 hpi in the presence of 9 μM cyclic peptomers or DMSO; a decrease in inclusion number or size suggests inhibition of binding, entry, EB-RB differentiation, or replication. Production of infectious progeny, which assays RB-EB re-differentation (including production of pre-packaged effectors) and/or release of EBs, was assayed by collecting EBs at 48 hpi, and infecting fresh monolayers for 24 hpi, and then quantifying inclusion formation. INP0400, a known T3SS inhibitor was used as a control (33). 4EpDN but not 4EpDN 2Sar decreased primary inclusion formation ~50% but inhibited formation of infectious progeny ~98%, without affecting host cell viability as assayed by LDH release (Fig. 9). Together, these results suggest that 4EpDN may predominantly inhibit production or secretion of pre-packaged *C. trachomatis* effectors rather than those made during RB replication.

**Figure 9.**
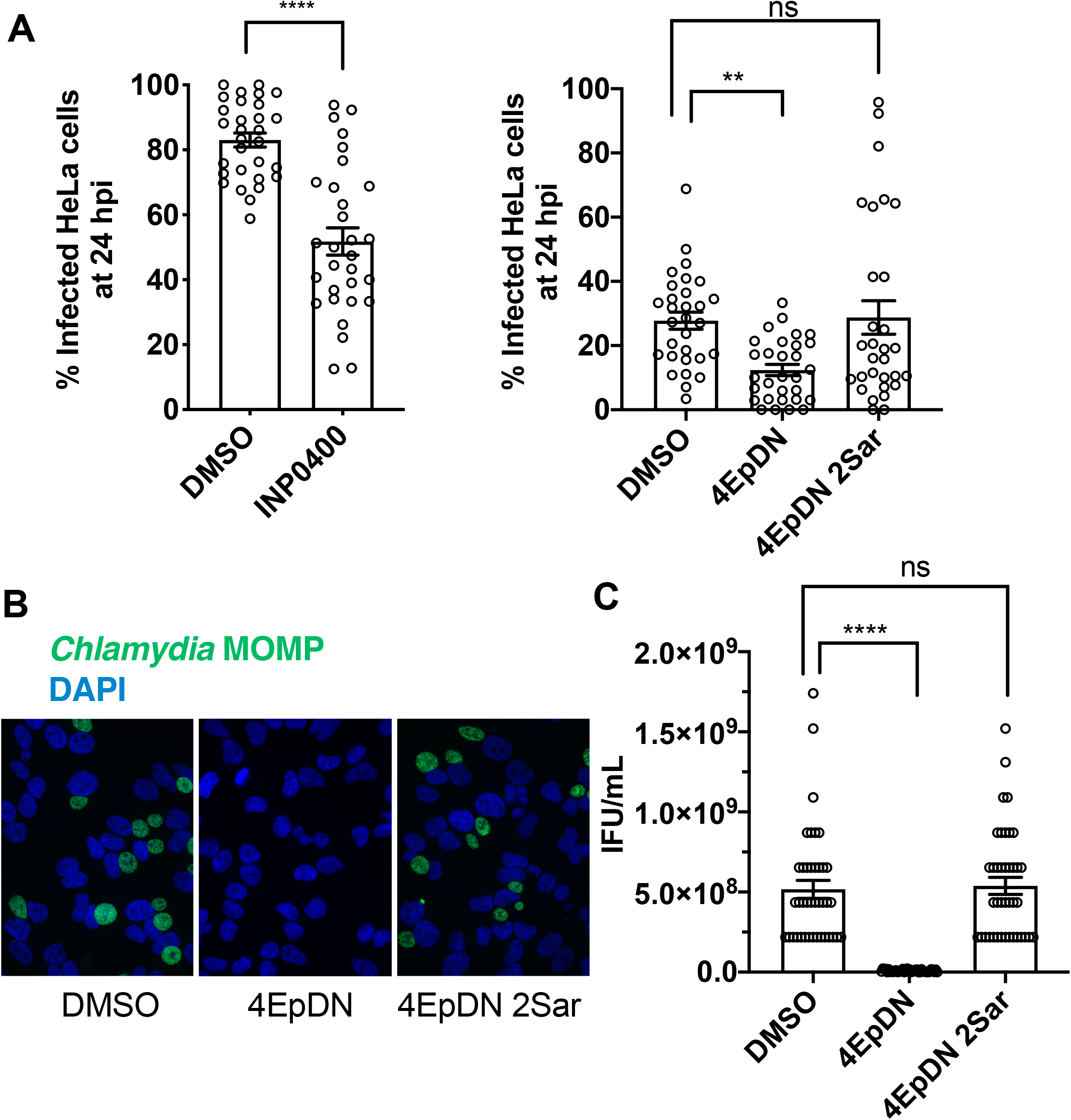
The cyclic peptomer 4EpDN inhibits *Chlamydia* infection. **(A)** HeLa cells were infected with *C. trachomatis* L2 at a multiplicity of infection (MOI) of three (left hand panel) or one (right hand panel) in the presence of 9 μM cyclic peptomers, 30 μM INP0400, or DMSO. Cells were stained for the *Chlamydia* major outer membrane protein (MOMP) and nucleic acids (DAPI), and imaged after 24 hrs of infection to determine the number of infected cells (primary infection). **(B-C)** Infectious elementary bodies (EB) were harvested after 48hrs of HeLa cell infection in the presence of inhibitors and used to infect fresh HeLa cells without applying inhibitors (secondary infection). After 24 hrs, cells were imaged as in **(A)**. Representative images (**B**) and infectious units/mL (**C**) are shown from three to four independent experiments. One-way ANOVA with Dunnett’s multiple-comparison test was used. **, P < 0.01, ****, P < 0.0001, ns: not significant.

## DISCUSSION

In this study we further developed cyclic peptomers as T3SS inhibitors and investigated their effects on various virulence mechanisms and pathogens. A newly synthesized cyclic peptomer derivative, 4EpDN, exhibited an IC_50_ in the low μM range. 4EpDN inhibits secretion through the T3SS of a number of pathogens including the nosocomial ESKAPE pathogen *Pseudomonas aeruginosa*, enteropathogenic *Yersinia*, *Salmonella* and the obligate intracellular pathogen *Chlamydia trachomatis*. 4EpDN does not inhibit secretion from two other secretion systems – the flagellar T3SS and the Tat secretion system. Cyclic peptomers do not inhibit T3SS gene expression, but affect locatization of the T3SS basal body component YscD, indicating disruption of normal T3SS assembly. These data suggest that cyclic peptomers specifically inhibit the injectisome T3SS in a broad spectrum of pathogens.

### Structure activity relationship analysis resulted in a T3SS inhibitor with an IC_50_ in the low μM range

Through alanine and stereochemistry scans, we identified 4EpDN, a cyclic peptomer with an IC_50_ of 4 μM, as inhibiting secretion of T3SS effector proteins in *P. aeruginosa* and 1 μM in inhibiting the *Salmonella* SPI-1 T3SS. Compared to previously published T3SS inhibitors (Table 2), this low μM activity is encouraging. The only published T3SS inhibitors with comparable IC_50_ are the phenoxyacetamides (MBX 2359 and its optimized derivatives), which inhibit *P. aeruginosa* T3SS secretion at 1-3 μM (28). Stereoisomers of 1EpDN showed a wide range of potencies, suggesting that differences in their three dimensional structures affect their biological activity. 9EpDN is a true enantiomer of 1EpDN with an IC_50_ of ~13 μM, a lower potency than the 1EpDN parental compound of ~8 μM. Importantly, the activity of these isomers do not positively correlate with solubility (Fig. S4), indicating that the observed activity is due to a specific molecular reaction rather than a non-specific biophysical effect due to aggregation. Furthermore, the presence of nonionic detergent did not affect activity of compounds (Fig. 4). Moreover, 4EpDN is potent at concentrations significantly lower than its solubility (Table S1). These data suggest that 4EpDN is an active cyclic peptomer with specific T3SS inhibitory activity.

**Table 2:**
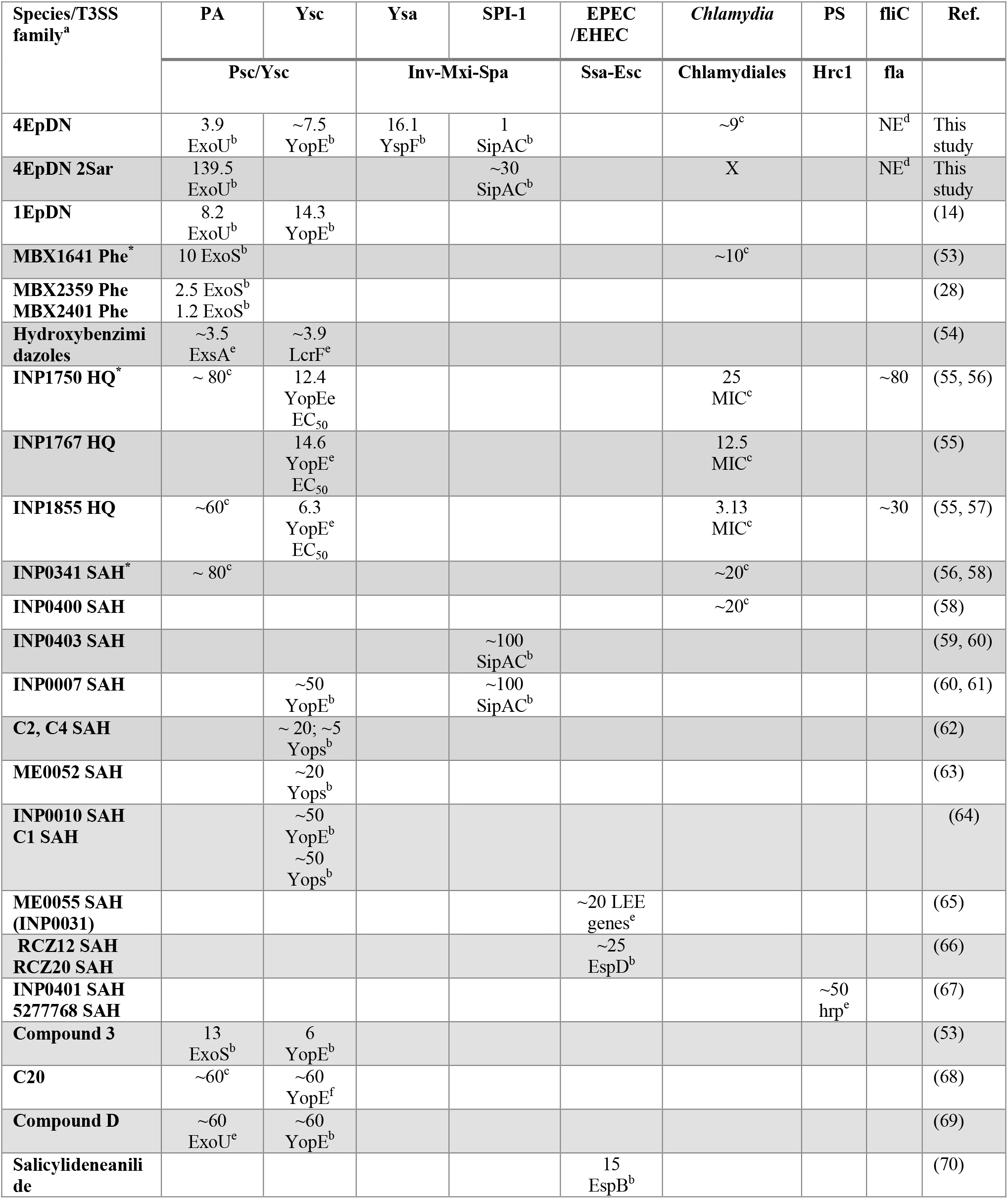
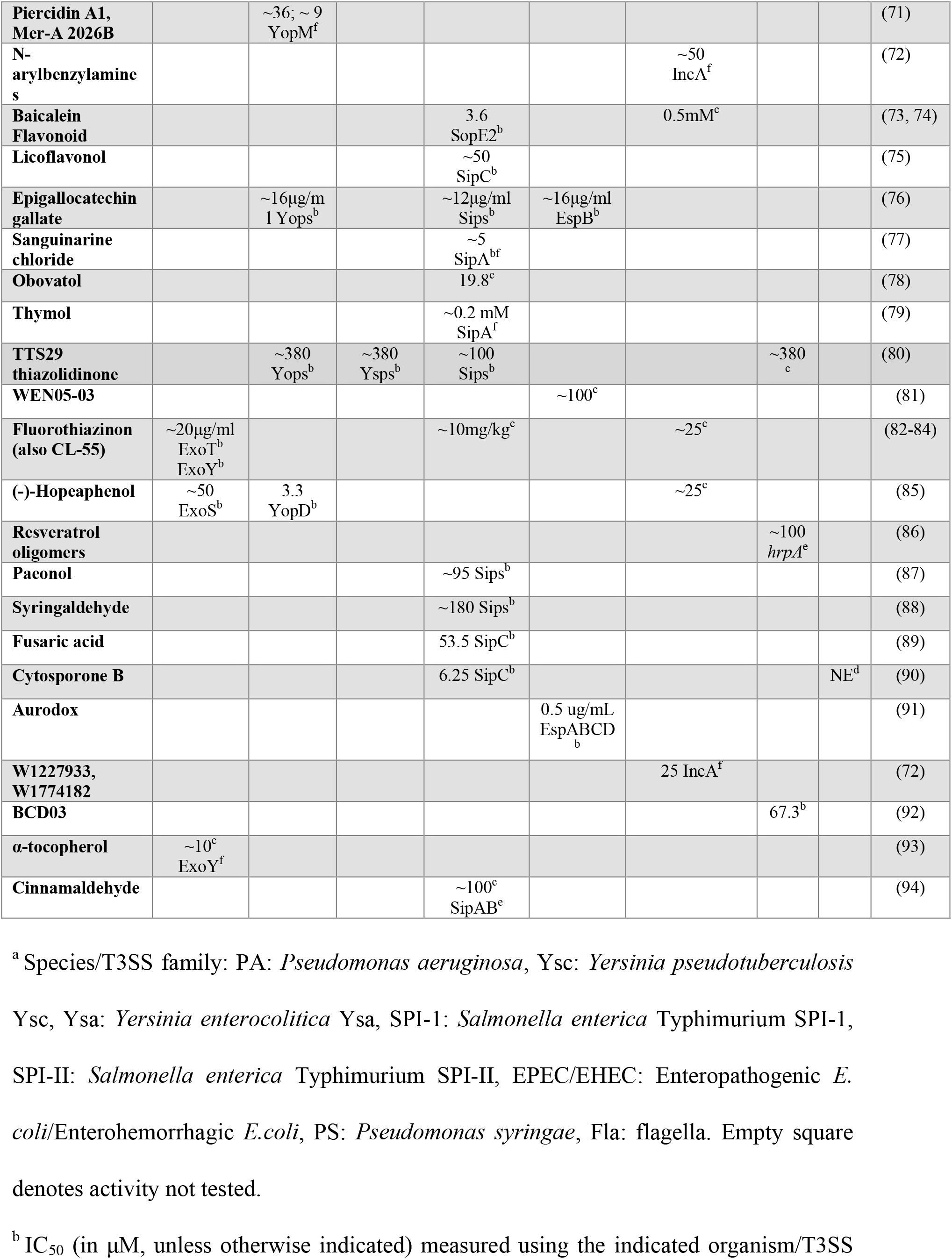
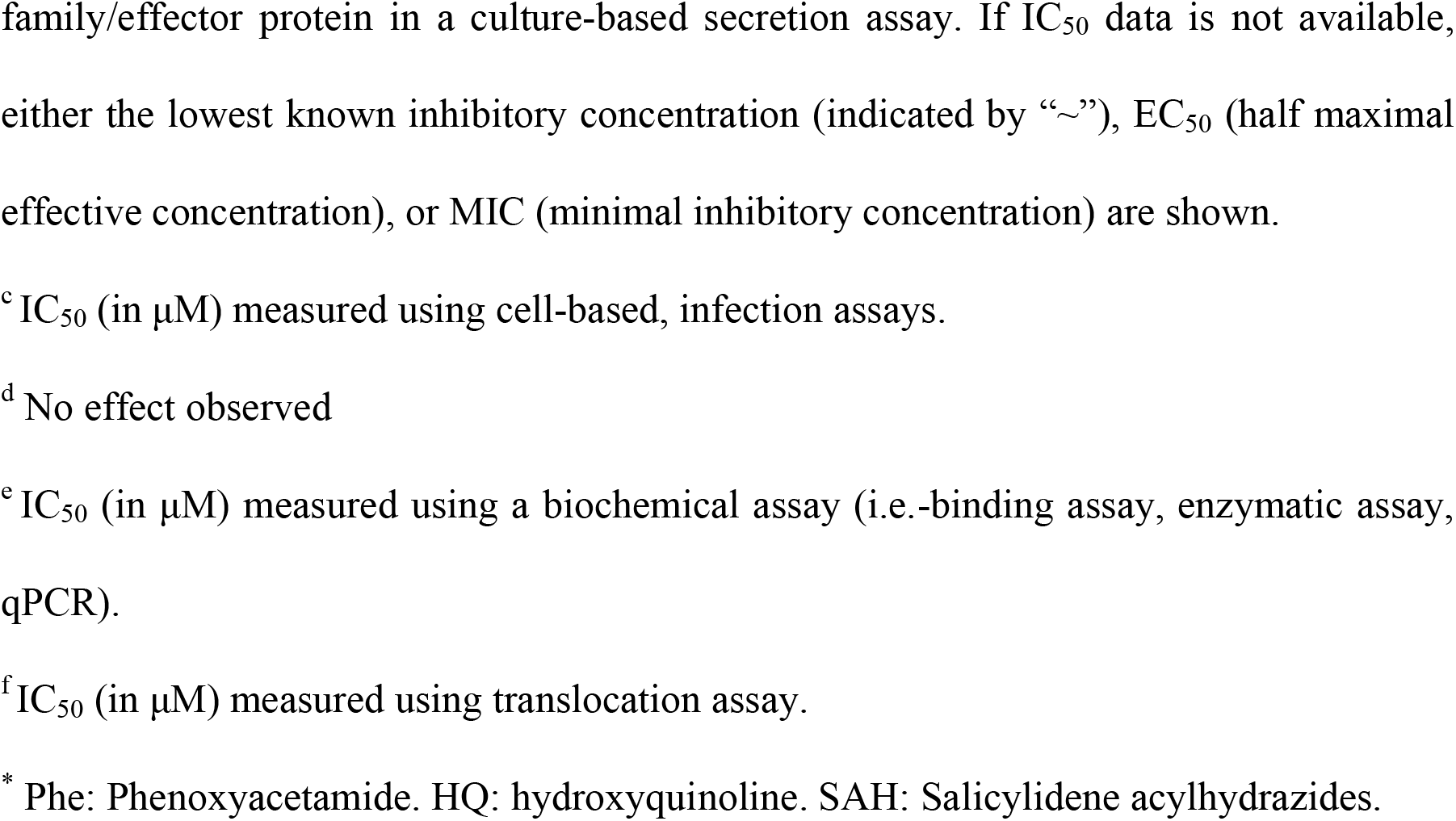
Efficacy of cyclic peptomers and other type III secretion system inhibitors.

### Cyclic peptomers act as broad-spectrum, but specific, inhibitors of the injectisome T3SS

Secretion of protein substrates through the injectisome T3SS, the flagellar system, and the Tat secretion system require the proton motive force (34–36). Although cyclic peptomers inhibited secretion from the injectisome T3SS, they did not inhibit the Tat system and only weakly inhibited flagellar substrate secretion, suggesting that the proton motive force is unaffected, as we previously suggested (14), and that the cyclic peptomers do not inhibit bacterial secretion in general.

The 4EpDN cyclic peptomer demonstrated efficacy against the T3SSs of *P. aeruginosa*, *Y. pseudotuberculosis*, *Y. enterocolitica*, *Salmonella enterica* Typhimurium, and *Chlamydia trachomatis*, with an IC_50_ in the range of 1 μM (for the *Salmonella* SPI-1 T3SS) to ~16 μM (for the *Y. pseudotuberculosis* Ysa T3SS) (Table 2). Based on phylogenetic analysis of core T3SS proteins, T3SSs group into seven T3SS families, five of which contain T3SSs from human pathogens (37). 4EpDN has efficacy against T3SSs from three of these T3SS families: the Ysc (Ysc and Psc), Inv-Mxi-Spa (SPI-1 and Ysa), and *Chlamydiales*. Interestingly, the flagellar ATPase from *E. coli* falls at the root of the phylogenetic tree (38), distinct from other T3SS families. As 4EpDN impacted secretion through the flagellar T3SS significantly less than through the injectisome T3SS in the same bacterial species and under the same culture and experimental conditions, we reason that the pathway targeted by cyclic peptomers is common to all injectisome T3SSs but absent from the flagellar system.

### Cyclic peptomers decrease cell envelope localization of the T3SS basal body protein YscD and inhibit secretion of T3SS effector proteins, but do not inhibit T3SS gene expression

The T3SS is a complex system made of ~20 different proteins and is assembled in a hierarchical manner prior to secretion of effector proteins (22, 39). Our study showed that 4EpDN inhibited secretion of T3SS effector proteins in *Yersinia*, *Pseudomonas*, and *Salmonella* and blocked *Chlamydial* growth, which requires translocation of T3SS effector proteins in human cells. Importantly, 4EpDN did not decrease expression of T3SS genes in *Pseudomonas* or *Salmonella* (Fig. 6, Fig. S5), suggesting that cyclic peptomers do not act at the level of T3SS gene expression. However, the T3SS of *Yersinia*, *Pseudomona*s, *Salmonella*, and *Chlamydia* share a number of orthologous basal body components that, if targeted, could lead to disruption of effector protein secretion. Interestingly, 4EpDN reduced localization of the inner membrane ring protein YscD to the *Yersinia* cell envelope (Fig 3), indicating a perturbation to T3SS assembly or stability.

Surprisingly, the effect of 4EpDN on the T3SS did not significantly impact secretion of the *Pseudomonas* regulator ExsE. In *P. aeruginosa,* ExsA is the key positive transcriptional regulator of genes encoding the T3SS machinery and substrates. ExsE leads to sequestration of ExsA, preventing it from inducing T3SS gene expression through a complex partner switching pathway (29, 40, 41). Under secreting conditions, ExsE, a T3SS substrate, is secreted, enabling release of ExsA and thereby allowing increased transcription of T3SS genes (29). However, we did not detect a decrease in ExsE secretion in the presence of cyclic peptomers, in contrast to the marked decrease in effector protein secretion. These results are consistent with the data showing that 4EpDN did not inhibit T3SS gene expression in *P. aeruginosa*, as ExsE mediates elevation of gene expression in response to secretion. It is not completely clear why the cyclic peptomers inhibited secretion of effector substrates and ExsE differently, although the small size of ExsE compared to other effectors such as ExoU (~8.7 kDa versus 74 kDa) may be a factor.

### Cyclic peptomers strongly inhibit *Chlamydia* primary and secondary infection

4EpDN strongly inhibited *Chlamydia* from infecting HeLa cells during primary infection, and furthermore prevented *Chlamydia* from infecting subsequent host cells (Fig. 9). This highlights the potential of cyclic peptomers to prevent the spread of *Chlamydia* infection. *Chlamydia* relies on its T3SS effector proteins to interact with host factors, such as the actin cytoskeleton, Golgi network, endoplasmic reticulum, and microtubule network, to mediate invasion and intracellular growth (42). It is possible that compounds that inhibit these host pathways could interfere with *Chlamydial* growth (43–45). However, microscopic analysis of many cellular structures in HeLa cells in the presence of 4EpDN did not show any gross changes to the actin cytoskeleton, Golgi network, endoplasmic reticulum, or microtubule network at the concentration used in our *Chlamydia* infection, 9 μM (Fig. S6). *C. trachomatis* infection may cause infertility in female patients and eye damage, in addition to lung infections (13). Antibiotics, such as β-lactam antibiotics, are a common way to treat *Chlamydia* infection, but the chance of recurrence is high (46, 47). Current vaccine development efforts are underway but multiple challenges remain (48). There is increased demand for drugs against *Chlamydia* due to antibiotic resistance (49). The strong efficacy of cyclic peptomers highlights their potential for development as an anti-*Chlamydial* drug.

## MATERIALS AND METHODS

### Bacterial strains and culture conditions

The bacterial strains and cell lines used in this study are listed in Table 3. All cultures were grown with shaking at 250 rpm unless otherwise noted. *Y. pseudotuberculosis* was grown in 2xYT (2x yeast extract and tryptone) at 26°C overnight. To induce the T3SS, the cultures were subcultured to an optical density at 600 nm (OD_600_) of 0.2 into low-calcium medium (2xYT with 20 mM sodium oxalate and 20 mM MgCl_2_). *Y. enterocolitica* was grown in BHI (brain heart infusion) medium at 26°C overnight. The Ysc T3SS in *Y. enterocolitica* was induced using low-calcium BHI (BHI with 20 mM sodium oxalate and 20 mM MgCl_2_). The Ysa T3SS was induced as described previously (50) using ‘L media’ (1% Tryptone, 0.5% Yeast extract) with 290 mM NaCl at 26°C. *P. aeruginosa* and *S. enterica* were grown in Luria-Bertani (LB) medium overnight at 37°C. For *P. aeruginosa*, the T3SS was induced using low-calcium medium (LB with 5 mM EGTA and 20 mM MgCl_2_). SPI-1 T3SS secretion was assessed after subculturing into fresh LB at 37°C unless noted otherwise. HeLa cells (ATCC) were cultured in Dulbecco’s modified Eagle’s medium (DMEM) with 10% fetal bovine serum (FBS). All cell lines were incubated at 37°C with 5% CO_2_.

**Table 3:**
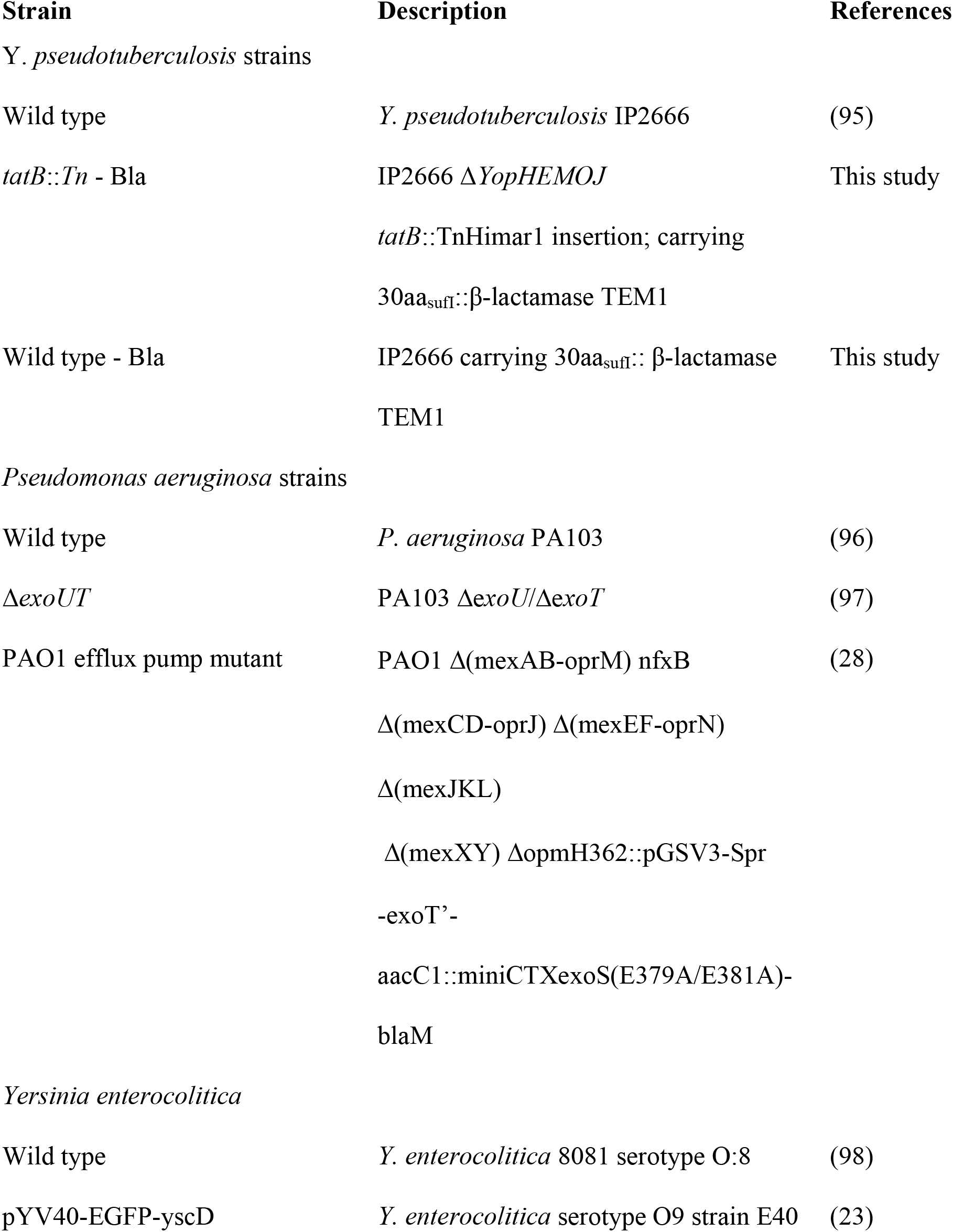
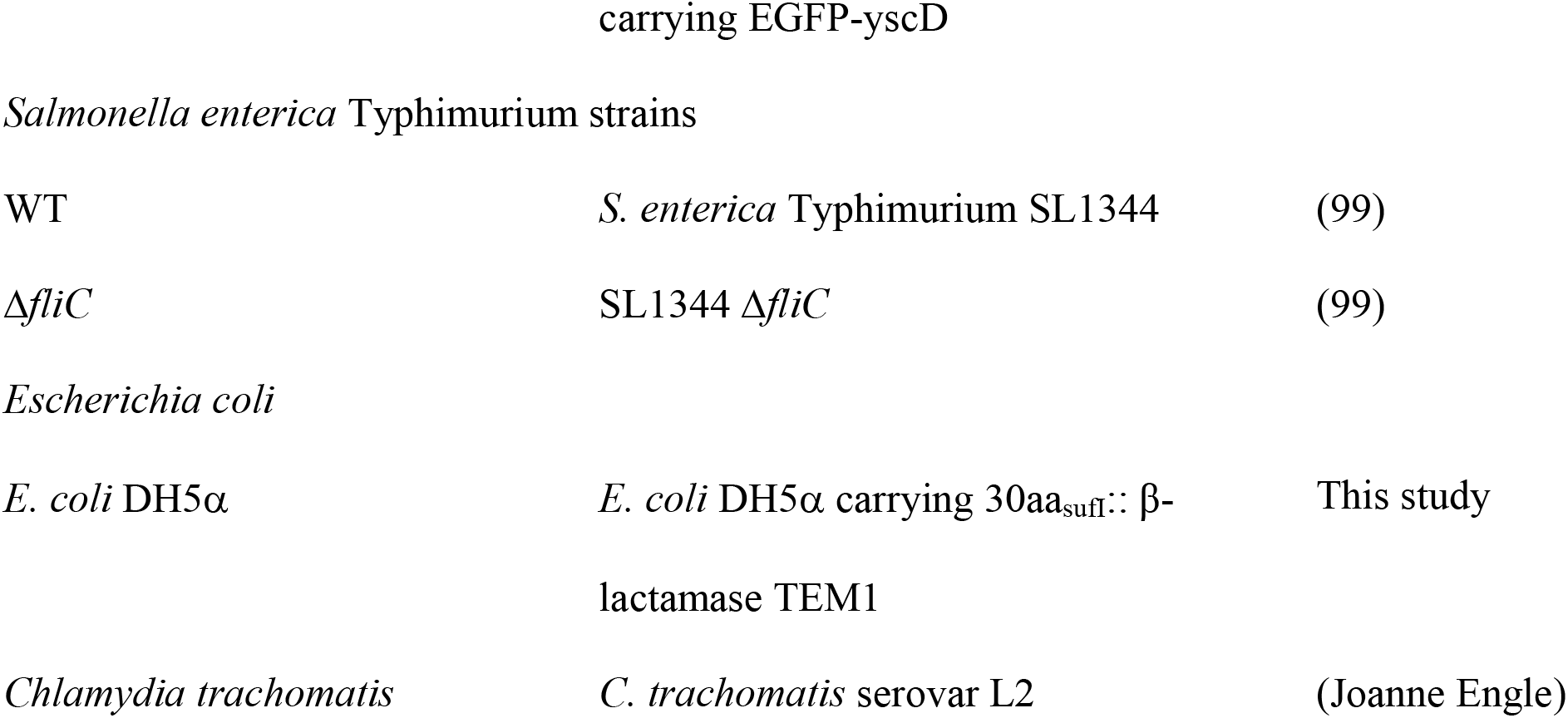
Bacterial strains used in this study.

### Preparation of bacteria for T3SS induction

Visualization of secreted proteins was carried out as described previously (25). Briefly, *Y. pseudotuberculosis*, *P. aeruginosa*, or *S. enterica* was grown in T3SS-inducing medium (as described above) in the presence of cyclic peptomers or an equivalent volume of DMSO at 37°C, 2 hrs for *Y. pseudotuberculosis* Ysc T3SS, 3 hrs for *P. aeruginosa*, 4 hrs for *S. enterica,* or at 26°C for 6 hrs for the *Y. enterocolitica* Ysa T3SS. The cultures were normalized to bacterial density (OD_600_) and then centrifuged for 15 min at 14,800 rpm. The supernatants were transferred to new tubes and mixed with trichloroacetic acid (TCA) to a final volume of 10% by vortexing vigorously for 30 s. Samples were incubated on ice for 1 hr and then spun down at 4°C for 15 min at 13,200 rpm. The supernatants were carefully removed, and pelleted proteins washed with acetone and spun down at 4°C for 15 min at 13,200 rpm for a total of three washes. The pellet was then resuspended in final sample buffer (FSB) and 20% dithiothreitol (DTT) and boiled for 15 min prior to SDS-PAGE. Tween-20 was added to the bacterial culture at the same time as the compounds in *S. enterica* secretion assays at 0.003% (v/v).

### T3SS secretion cargo quantification

Image Lab software (Bio-Rad) was used to quantify T3SS cargo protein bands relative to those of DMSO-treated controls. The WT *Y. pseudotuberculosis* YopE, *P. aeruginosa* ExoU, or *S. enterica* SipA, SipC, FliC and FliD bands in DMSO control samples were set to 1.00. To evaluate type III secretion of ExoS and ExsE in *P. aeruginosa*, Western blots against T3SS cargo were carried out, using a PVDF membrane (Millipore), blocking in 5% non-fat milk for 2 hrs at room temperature, and incubated at 4°C overnight with gentle shaking. Blots were washed three times for 5 minutes each in Phosphate-Buffered Saline with 0.1% Tween® 20 (PBST). Horseradish Peroxidase conjugated secondary antibody was then incubated for 1 hr at room temperature. Signals were detected with Luminol Kit after washing. ExoS-Bla, ExsE and SipC was visualized using β-lactamase (MA120370, Fisher Scientific) (7.5% gel), anti-ExsE antibody courtesy of Timothy Yahr (20% gel) and anti-SipC (ABIN335178, antibodies-online Inc.)(10% gel), respectively.

### YscD visualization assay

*Y. enterocolitica* expressing YscD-EGFP was cultured overnight in BHI supplemented with nalidixic acid (35 μg/mL) and diaminopimelic acid (50 μg/mL), at 26°C with shaking (23), followed by subculture into low calcium BHI medium (20mM NaOX, 20mM MgCl_2_, 0.4% Glycerol) with nalidixic acid and diaminopimelic acid to OD_600_ of 0.2 for 1.5 hrs. Compounds or an equivalent volume of DMSO was added prior to inducing the T3SS. After 3 hrs at 37°C with shaking, cells were pelleted and resuspended in PBS, spotted onto a 0.1% agarose pad, and imaged live at 63X/1.4 oil magnification using a Zeiss AxioImager widefield microscope. Analysis of YscD puncta was carried out in Imaris 8 using spot tracking analysis with the same arbitrary threshold to call bacterial cell and puncta for all samples.

### mRNA quantification

Overnight *P. aeruginosa* (PA103 or PA01) cultures were subcultured and shifted to T3SS inducing conditions (see above) in the presence of 60 μM 1EpDN, 60 μM 1EpDN 2Sar, or 50 μM MBX1641. Samples were taken after 3 hrs of induction. Overnight *Salmonella* cultures were subcultured into fresh LB with 0.3 M NaCl at 37°C in the presence of 9 μM 4EpDN, 4EpDN 2Sar, or equivalent DMSO. Samples were taken after 2 hrs and 4 hrs of induction. Samples were stored in RNAprotect reagent (Qiagen) and processed within a week. Total RNA was isolated using an RNAeasy Kit (Qiagen), according to the manufacturer’s instructions, followed by two rounds of Turbo DNAse (ThermoFisher scientific) treatment. A total of 2 μg of RNA was used to make cDNA and qPCR reactions were run with SYBR Green PCR master mix (Applied Biosystems). DNA helicase (*dnaB*) and 16S rRNA genes were used as a reference for *P. aeruginosa* and *Salmonella* samples, respectively. Two to three technical replicates were averaged for each sample. Primers used are listed in Table S2. Results were analyzed using Bio-Rad CFX software.

### Tat translocation assay

To make Tat targeting constructs, plasmid pMMB67EH (ATCC 37622) was digested with KpnI. TEM1 of β-latacmase was PCR amplified from *yopH*-Bla (courtesy of Melanie Marketon) using primers oHL210 and oHL217 (Table S2) and *sufI* signal peptide DNA was PCR amplified from genomic DNA of *Y. pseudotuberculosis* with primers oHL218 and oHL219 (Table S2). The digested pMMB67EH, TEM1 and *sufI* signal peptide DNA were assembled into a plasmid (*sufI*-Bla) using Gibson assembly.

WT *Yersinia* or *tatB::Tn* carrying *sufI*-Bla was grown in 2 x YT supplemented with 15μg/mL gentamicin at 26°C with shaking. Overnight cultures were subcultured to OD_600_ of 0.1 and grown for 1.5hr at 26°C with shaking. 5 mM IPTG was added to the culture for 0.5 hrs to allow for expression and translocation of SufI-Bla. Penicillin G (25μg/mL) was added to the cultures. Cultures were then treated with cyclic peptomers or DMSO and OD_600_ measured every hour up to 8hrs.

### *Chlamydia* infection and imaging

Primary Infections: HeLa cell monolayers were infected with *C. trachomatis*, Serovar L2 at an MOI of one in the presence of one of the following compounds at 9 μM: (a) DMSO, (b) 4EpDN, or (c) 4EpDN 2Sar. Cells were incubated for 24 hrs in the presence of the above compounds at 37°C, and then fixed with 4% Paraformaldehyde (PFA). Cells were stained for IncA (*Chlamydia* inclusion membrane marker), DNA, and MOMP (*Chlamydi*a major outer membrane protein). The percentage of cells infected (i.e. stained positively for the listed *Chlamydia* markers) in the presence of the compounds was quantified using confocal microscopy. 10 randomly selected fields of view were measured per experiment. Data represents three biological replicates.

Secondary Infections: HeLa cell monolayers were infected with *C. trachomatis*, Serovar L2 at an MOI of 1.0 in the presence of one of the following compounds at 9 μM: (a) DMSO, (b) 4EpDN, or (c) 4EpDN 2Sar. Cells were incubated for 48 hrs in the presence of the above compounds at 37°C. Infected cells were then lysed, and the lysate was applied to fresh HeLa monolayers to enumerate infectious particles. These secondary infections were fixed in 4% PFA at 24 hpi, and were stained against MOMP and DNA. Infectious units per mL (IFU/mL) were calculated by averaging the number of infected cells in each of 10 randomly selected fields of view at 60X magnification on a confocal microscope, and multiplying this by the appropriate dilution and area factors. Data represents four biological replicates.

### Cytological Profiling (CP)

Briefly, HeLa cells were cultured and seeded into 384-well at 2,500 cells/well. After 48 hrs, compounds were added using a Janus MDT robot (PerkinElmer). Two stain sets were used; Stain set 1: Hoechst, EdUrhodamine, anti-Phosphohistone H3, and GM130, Stain Set 2: Hoechst, FITC-alpha tubulin, rhodamine-phalloidin, and Calnexin. For stain set 1, cells were incubated with 20 μM EdUrhodamine for 1 hr prior to fixing in 4% formaldehyde solution in PBS for 20 min. Cells were then washed with PBS and permeabilized with 0.5 % Triton-X in PBS for 10 min before blocking with 2 % BSA in PBS solution for at least 1 hr. Following this, cells were incubated with primary antibodies overnight at 4°C. The following day, excess primary antibody was washed off with PBS and Alexa-488 and Alexa-647 secondary antibodies and Hoechst solution were incubated for 1 hr. Plates were washed with PBS and preserved with 0.1% sodium azide in PBS solution prior to imaging. For stain set 2, cells were fixed with a 4 % formaldehyde solution in PBS for 20 min. Cells were then washed with PBS and permeabilized with 0.5 % Triton-X in PBS for 10 min before blocking with 2 % BSA in PBS solution for at least 1 hr. Following this, cells were incubated with primary antibodies overnight at 4°C. After blocking, the cells were washed, and incubated with FITC conjugated anti-alpha tubulin antibody and rhodamine-phalloidin overnight at 4°C. The following day the cells were washed and then incubated with secondary Alexa-647 and Hoechst stain for 1 hr. Plates were washed with PBS and preserved with 0.1% sodium azide in PBS solution prior to imaging.

Two images per well were captured with an ImageXpress Micro XLS automated epiflourescent microscope (Molecular Devices, Sunnyvale). Images were then processed as described (51). Briefly, initial image processing was performed using MetaXpress image analysis software, using built-in morphometry metrics, the multiwavelength cell scoring, transfluor, and micronuclei modules. Custom written scripts were used to compare the treated samples with the DMSO control wells, and then to convert each feature to a “histogram difference” (HD) score. This produced a 452-feature vector CP fingerprint. Compound treatment wells were labeled as dead if the cell count for the treatment well was < 10% of the median cell count in the treatment plate. In addition to the CP fingerprint, feature cell counts (nuclei, EdU S-phase, and phospho-histone H3) were used to determine effects of compounds on HeLa cell replication.

#### Synthesis of cyclic peptomers

### Cyclic peptide synthesis

Peptides were synthesized using standard Fmoc solid-phase peptide synthesis, utilizing the submonomer approach for peptoid synthesis (52), either at room temperature or with microwave assistance. Cyclization was done in solution at a high dilution. Fmoc-Xaa (10 mmol) was added to a flame-dried round-bottomed flask and dried in a vacuum desiccator with phosphorous pentoxide overnight. 50 ml of dry dichloromethane (DCM) was cannula transferred into the flask, followed by 2.5 ml of N,Ndiisopropylethylamine (DIPEA) transferred via syringe. After sonication for 10 min, 5g of 2-chlorotrityl resin was added under a stream of nitrogen and allowed to shake for 4 hrs. The resin was capped with a 15 ml solution of 1:2:17 methanol (MeOH):DIPEA:dimethylformamide (DMF) (3 times for 15 min each). The resin was washed with DMF (3 times with 15 ml each) followed by DCM (3 times with 15 ml each). The loading value was calculated by determining the mass increase of dried, loaded resin.

### Amino acid coupling at room temperature

Four equivalents (eq) of Fmoc-Xaa, 8 eq of DIPEA, and 4 eq of 1-[Bis(dimethylamino)methylene]-1H-1,2,3-triazolo[4,5-b]pyridinium 3-oxide hexafluorophosphate, **H**exafluorophosphate **A**zabenzotriazole **T**etramethyl **U**ronium (HATU) were added to the resin in DMF. The reaction mixture was agitated via shaking for 45 min and then drained. The resin was washed with DMF (3 times with 3 ml each) and DCM (3 times with 3 ml each). The reaction was monitored by liquid chromatography-mass spectrometry (LC-MS) and repeated until the starting material was no longer observed. For microwave conditions, a solution of 4 eq of Fmoc-Xaa, 4 eq of HATU, and 6 eq of DIPEA in DMF was allowed to prereact for 5 min. This solution was added to the deprotected peptide on-resin and allowed to react for 10 min at 50°C under microwave heating. The solution was drained, and the resin was washed with DMF (3 times with 3 ml each) and DCM (3 times with 3 ml each). The reaction was monitored by LC-MS and repeated until the starting material was no longer observed.

### Coupling of BrAcOH at room temperature

A solution of 10 eq of bromoacetic acid (BrAcOH) and 5 eq of N,N=-diisopropylcarbodiimide (DIC) in DMF was allowed to prereact for 10 min. This solution was added to the deprotected peptide on-resin. The reaction mixture was agitated via shaking for 45 min and then drained. The resin was washed with DMF (3 times with 3 ml each) and DCM (3 times with 3 ml each). The reaction was monitored by LC-MS and repeated until the starting material was no longer observed. The reaction was monitored by LC-MS and repeated until the starting material was no longer observed.

### Peptoid side chain addition

A solution of 5 eq of the desired amine was prepared in a minimum volume of DMF. The resin containing the BrAc-peptide was swollen with DCM for 5 min prior to reaction. The amine was added, and the reaction mixture was agitated via shaking for 3 to 20 hrs. The solution was drained, and the resin was washed with DMF (3 times with 3 ml each) and DCM (3 times with 3 ml each). The reaction was monitored by LC-MS and repeated until the starting material was no longer observed.

### Removal of the N-Fmoc protection group at room temperature

A solution of 2% piperidine and 2% 1,8-diazabicyclo[5.4.0]undec-7-ene (DBU) in DMF was added to the resin. The reaction mixture was agitated via shaking for 20 min and then drained. The resin was washed with DMF (3 times with 3 ml each) and DCM (3 times with 3 ml each). For microwave conditions, a solution of 2% piperidine and 2% DBU in DMF was added to the resin. The reaction mixture was allowed to react for 5 min at 50°C under microwave heating and then drained. The resin was washed with DMF (3 times with 3 ml each) and DCM (3 times with 3 ml each).

### Peptide cleavage

Complete linear peptides were cleaved off the resin in 5 resin volumes of 2.5% trifluoroacetic acid (TFA) in DCM for 4 min, three times, with a 5-resin-volume DCM wash between steps. Solvent was removed under N2, followed by dissolution in acetone or DCM and evaporation under reduced pressure. Residual TFA was removed in vacuo overnight.

### Cyclization with COMU

Linear peptides were dissolved in 20 ml of dry acetonitrile (ACN) with 4 eq of DIPEA and added dropwise (final concentration, 1 mg crude peptide per ml) to a solution of 1:1 tetrahydrofuran (THF)-ACN containing 2 eq of (1-cyano-2 ethoxy-2 oxoethylidenaminooxy) dimethylamino-morpholinocarbenium hexafluorophosphate (COMU). Reaction mixtures were stirred for 0.5 to 24 hrs, until complete cyclization was achieved as monitored by LC-MS. The reaction mixture was reduced in vacuo for purification via high pressure liquid chromatography (HPLC).

### Purification of peptides

COMU by-products were removed after solution-phase cyclization on a Biotage Isolera Prime system equipped with a SNAP Ultra-C18 30g column eluting with H2O-acetonitrile modified with 0.1% TFA. The mass spectra of all peptides are shown in Fig. S7 in the supplemental material.

### Proton NMR of peptides

Peptides were analyzed through nuclear magnetic resonance (NMR) spectroscopy measured in ppm and were obtained on a 500 MHz spectrometer using CDCl_3_ (δ=7.26) as an internal standard for ^1^H NMR. Identity of compounds for SAR study was confirmed by LCMS and ^1^H-NMR (Fig. S7).

### Kinetic Solubility

A 15 mM stock of the compounds in DMSO was prepared. 125uL of M9 and DMEM (no antibiotics) was dispensed into 96 v-bottom plate. One microliter of 15mM stock compound was added to make a solution of 120μM final concentration with 0.8% DMSO. The solution was shaken at 37°C for ~2 hrs. The solution was passed over a 0.7 μM glass fiber filter. Then the solution was diluted 1:4 in acetonitrile to crash out any proteins. The solution was centrifuged at 500 x g for 10 minutes. Avoiding the pellet, 10 μL of supernatant was added to a fresh plate with 90 μL of acetonitrile. The final dilution is 40 times lower. 10μL of 40x dilution of solution was injected on the Orbi-trap. A 1μM standard was used for the ratiometric comparison and the assay was done in triplicate.

### Cyclic peptide manipulation

Stock peptides were stored at 15 mM at −70°C and were prediluted in DMSO prior to experiments. All treatment and control pairs, in all assays, had the same DMSO volumes.

### Statistical analysis

GraphPad Prism 8 (GraphPad Software, La Jolla, CA, USA) was used to calculate the mean, standard error of the mean, median, standard error of median, and one-way ANOVA values shown.

## ACKNOWLEDGEMENTS

The authors acknowledge the National Institutes of Health grant R01AI141511 (to V.A. and R.S.L.) and K99AI139281 (to H.L.) for support. We thank Benjamin Abrams (University of California, Santa Cruz) for technical support on the YscD spot tracking analysis. We thank Timothy Yahr for the anti-ExsE antibody.

---

**Table S1:**
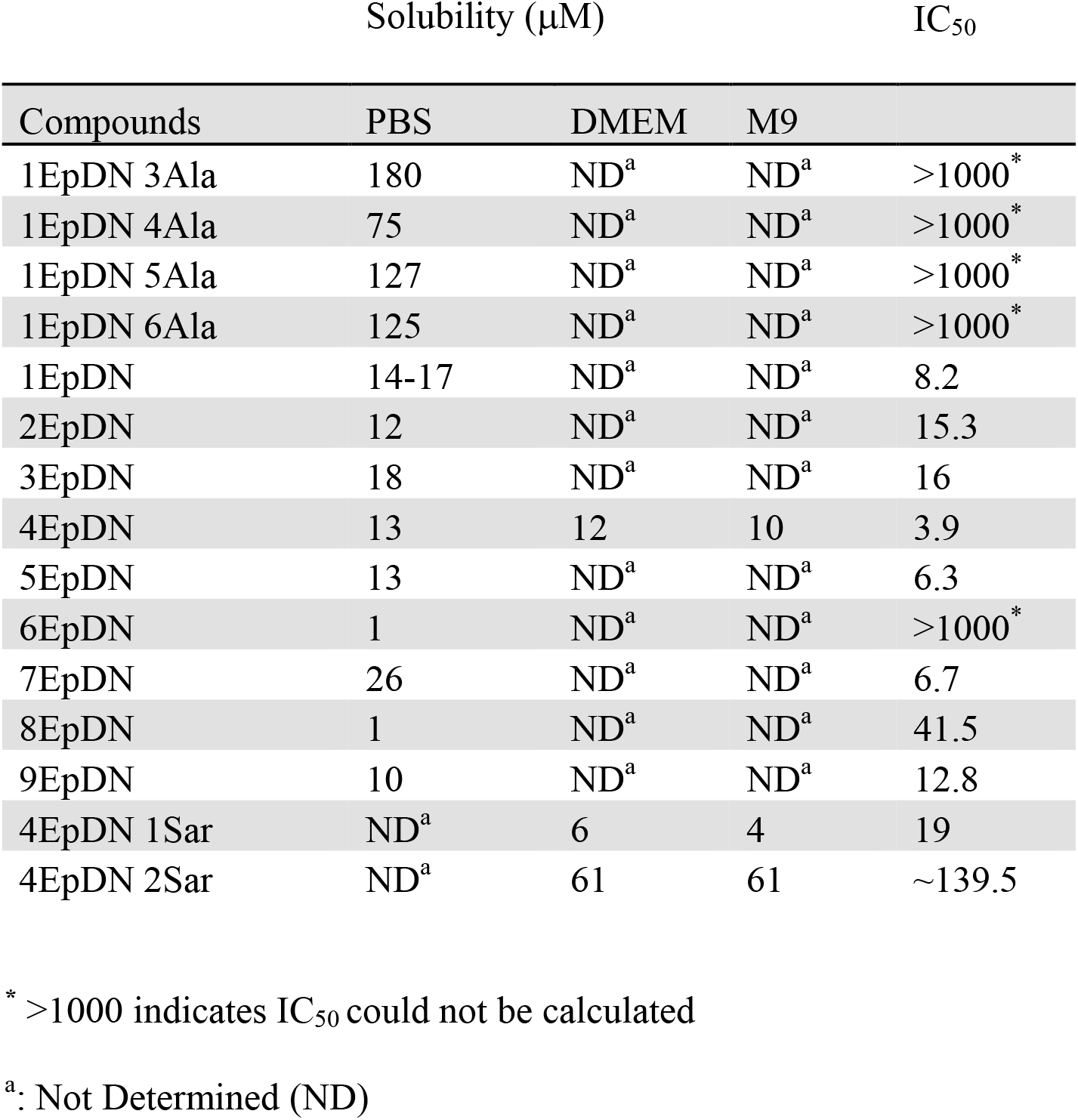
Solubility of cyclic peptomers.

**Table S2.**
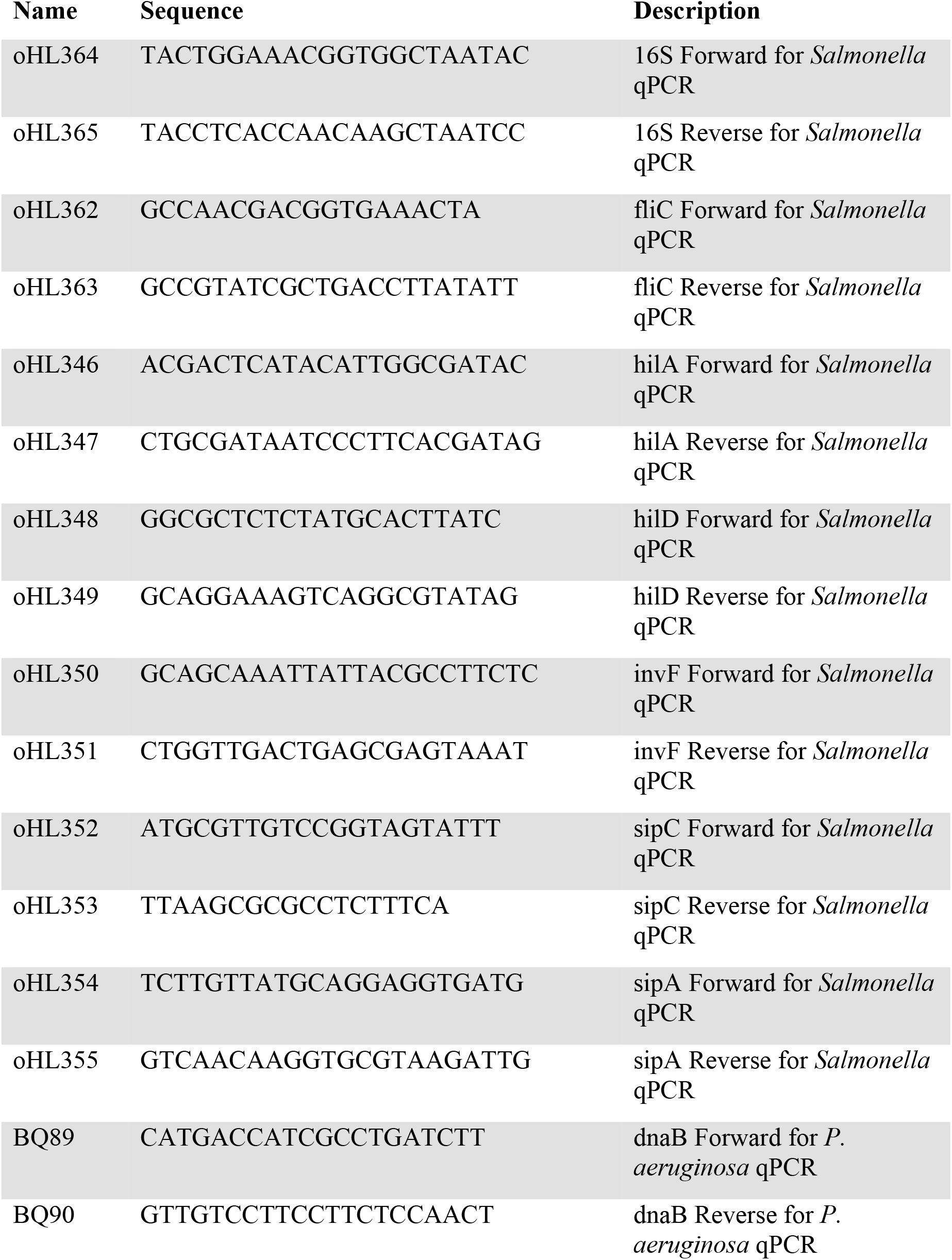
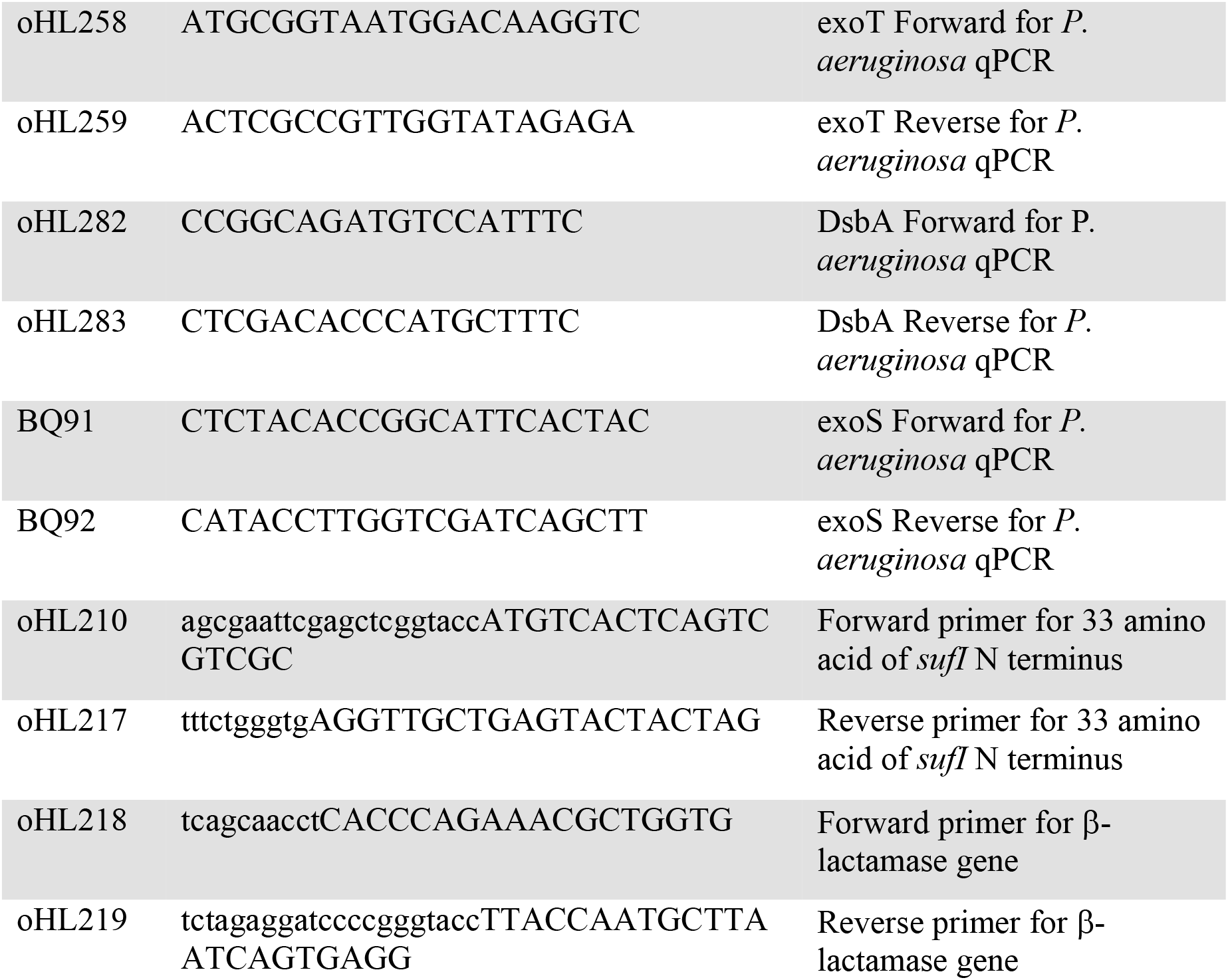
Primers used in this study.

## Figure legends

**Figure S1: 1EpDN alanine/Sarcosine scan suggests peptoid sidechains are important for biological activity. (A)** Structures of 1EpDN alanine derivatives. D-form of side chain is shown in red. **(B)** WT *P. aeruginosa* PA103 was grown under T3SS-inducing conditions with increasing concentrations of cyclic peptomers. Secretion of T3SS cargo into the culture supernatant was assessed by precipitating secreted proteins and visualizing them with Coomassie blue. ExoU band intensities were quantified and normalized to that of the DMSO control. The results are from at least two independent experiments. Nonlinear curve fitting is shown to depict the trend of inhibition.

**Figure S2: Secretion of *Salmonella* T3SS substrate in the presence of non-ionic detergents. (A)** *Salmonella enterica* Typhimurium was grown in LB with increasing concentrations of NP-40, Tween 20, or Triton X-100. Secretion of SPI-1 T3SS effector SipC into the culture supernatant was assessed by precipitating secreted proteins and visualizing them with Coomassie blue. **(B)** Secretion of SipC in the presence of increasing concentrations of Tween 20 or Triton X-100 was detected by Western Blot. 0.003% Tween-20 is the highest concentration of Tween-20 that resulted in little effect on secretion of SipC.

**Figure S3: Cyclic peptomers do not affect secretion of flagellar proteins.** *Salmonella enterica* Typhimurium was grown in LB with increasing concentrations of cyclic peptomers. Secretion of flagellar structural proteins FliC and FliD were assessed by precipitating the secreted proteins and visualizing them with Coomassie blue. A Δ*fliC* mutant and azithromycin (100), which inhibits flagellin secretion, were used as controls. The SPI-1 mutant and WT *Salmonella* were both tested, as flagella substrates can be secreted through both flagellar and SPI-1 T3SS systems.

**Figure S4**: **Relationship between solubility and activity of cyclic peptomers**. IC_50_ of stereoisomers and their solubility (table S1) were plotted on a log_10_ scale. Average solubility was used when the solubility was measured in different conditions.

**Figure S5: Cyclic peptomers do not affect transcription of T3SS genes in *Salmonella*.** *Salmonella enterica* Typhimurium was grown in LB with 300 mM NaCl in the presence of 9 μM cyclic peptomers or DMSO. Samples were taken 2 hrs (**A**) and 4 hrs (**B**) after addition of compounds at 37°C and expression of flagellar (*fliC*) and injectisome T3SS (*hilA, hilD, invF, sipC, sipA*) genes were assessed using qPCR. Data are from two replicates, analyzed by one-way ANOVA showing no significant difference between control and treatments.

**Figure S6: Effect of cyclic peptomers on HeLa cells.** HeLa cells were incubated with compounds for 48 hrs. Cells were then stained with: **(A)** Stain Set 1-Hoechst, FITC-alpha tubulin, rhodamine-phalloidin (actin), and Calnexin (ER induced protein); or **(B)** Stain set 2-Hoechst, EdU-rhodamine (S-phase detection), anti-Phosphohistone H3 (mitosis marker), and GM130 (Golgi matrix protein). Representative images of cells treated with different concentrations of 4EpDN or DMSO are shown. **(C)** Quantification of all cell features for 4EpDN-treated cells. The total CP score is the square root of sum of square of the difference between treatment and DMSO for all measured features.

**Figure S7:** Characterization of cyclic peptomers. Drawn structures, SMILE structures, molecular weight, LCMS Spectra, and ^1^H-NMR Spectra are shown.

## REFERENCES

1. Santajit S, Indrawattana N. 2016. Mechanisms of Antimicrobial Resistance in ESKAPE Pathogens. Biomed Res Int 2016:2475067.

2. Ma YX, Wang CY, Li YY, Li J, Wan QQ, Chen JH, Tay FR, Niu LN. 2020. Considerations and Caveats in Combating ESKAPE Pathogens against Nosocomial Infections. Adv Sci (Weinh) 7:1901872.

3. Duncan MC, Linington RG, Auerbuch V. 2012. Chemical inhibitors of the type three secretion system: disarming bacterial pathogens. Antimicrob Agents Chemother 56:5433–41.

4. Calvert MB, Jumde VR, Titz A. 2018. Pathoblockers or antivirulence drugs as a new option for the treatment of bacterial infections. Beilstein J Org Chem 14:2607–2617.

5. Relman DA, Lipsitch M. 2018. Microbiome as a tool and a target in the effort to address antimicrobial resistance. Proc Natl Acad Sci U S A 115:12902–12910.

6. Mohajeri MH, Brummer RJM, Rastall RA, Weersma RK, Harmsen HJM, Faas M, Eggersdorfer M. 2018. The role of the microbiome for human health: from basic science to clinical applications. Eur J Nutr 57:1–14.

7. Abby SS, Rocha EP. 2012. The non-flagellar type III secretion system evolved from the bacterial flagellum and diversified into host-cell adapted systems. PLoS Genet 8:e1002983.

8. Diepold A, Armitage JP. 2015. Type III secretion systems: the bacterial flagellum and the injectisome. Philos Trans R Soc Lond B Biol Sci 370.

9. Diepold A, Wagner S. 2014. Assembly of the bacterial type III secretion machinery. FEMS Microbiol Rev 38:802–22.

10. Fasciano AC, Shaban L, Mecsas J. 2019. Promises and Challenges of the Type Three Secretion System Injectisome as an Antivirulence Target. EcoSal Plus 8.

11. Tabor DE, Oganesyan V, Keller AE, Yu L, McLaughlin RE, Song E, Warrener P, Rosenthal K, Esser M, Qi Y, Ruzin A, Stover CK, DiGiandomenico A. 2018. Pseudomonas aeruginosa PcrV and Psl, the Molecular Targets of Bispecific Antibody MEDI3902, Are Conserved Among Diverse Global Clinical Isolates. J Infect Dis 218:1983–1994.

12. DiGiandomenico A, Keller AE, Gao C, Rainey GJ, Warrener P, Camara MM, Bonnell J, Fleming R, Bezabeh B, Dimasi N, Sellman BR, Hilliard J, Guenther CM, Datta V, Zhao W, Gao C, Yu XQ, Suzich JA, Stover CK. 2014. A multifunctional bispecific antibody protects against Pseudomonas aeruginosa. Sci Transl Med 6:262ra155.

13. Ha**cker G. 2018. Biology of chlamydia. Springer, Cham, Switzerland.

14. Lam H, Schwochert J, Lao Y, Lau T, Lloyd C, Luu J, Kooner O, Morgan J, Lokey S, Auerbuch V. 2017. Synthetic cyclic peptomers as type III secretion system inhibitors. Antimicrob Agents Chemother doi:10.1128/AAC.00060-17.

15. Gophna U, Ron EZ, Graur D. 2003. Bacterial type III secretion systems are ancient and evolved by multiple horizontal-transfer events. Gene 312:151–63.

16. Bent ZW, Branda SS, Young GM. 2013. The Yersinia enterocolitica Ysa type III secretion system is expressed during infections both in vitro and in vivo. Microbiologyopen 2:962–75.

17. Venecia K, Young GM. 2005. Environmental regulation and virulence attributes of the Ysa type III secretion system of Yersinia enterocolitica biovar 1B. Infect Immun 73:5961–77.

18. Young BM, Young GM. 2002. Evidence for targeting of Yop effectors by the chromosomally encoded Ysa type III secretion system of Yersinia enterocolitica. J Bacteriol 184:5563–71.

19. Hallstrom KN, McCormick BA. 2016. The type three secreted effector SipC regulates the trafficking of PERP during Salmonella infection. Gut Microbes 7:136–45.

20. Singh PK, Kapoor A, Lomash RM, Kumar K, Kamerkar SC, Pucadyil TJ, Mukhopadhyay A. 2018. Salmonella SipA mimics a cognate SNARE for host Syntaxin8 to promote fusion with early endosomes. J Cell Biol 217:4199–4214.

21. Lilic M, Galkin VE, Orlova A, VanLoock MS, Egelman EH, Stebbins CE. 2003. Salmonella SipA polymerizes actin by stapling filaments with nonglobular protein arms. Science 301:1918–21.

22. Deng W, Marshall NC, Rowland JL, McCoy JM, Worrall LJ, Santos AS, Strynadka NCJ, Finlay BB. 2017. Assembly, structure, function and regulation of type III secretion systems. Nat Rev Microbiol 15:323–337.

23. Diepold A, Amstutz M, Abel S, Sorg I, Jenal U, Cornelis GR. 2010. Deciphering the assembly of the Yersinia type III secretion injectisome. EMBO J 29:1928–40.

24. Diepold A. 2019. Assembly and Post-assembly Turnover and Dynamics in the Type III Secretion System. Curr Top Microbiol Immunol doi:10.1007/82_2019_164.

25. Diepold A, Sezgin E, Huseyin M, Mortimer T, Eggeling C, Armitage JP. 2017. A dynamic and adaptive network of cytosolic interactions governs protein export by the T3SS injectisome. Nat Commun 8:15940.

26. Kudryashev M, Stenta M, Schmelz S, Amstutz M, Wiesand U, Castano-Diez D, Degiacomi MT, Munnich S, Bleck CK, Kowal J, Diepold A, Heinz DW, Dal Peraro M, Cornelis GR, Stahlberg H. 2013. In situ structural analysis of the Yersinia enterocolitica injectisome. Elife 2:e00792.

27. Ha UH, Wang Y, Jin S. 2003. DsbA of Pseudomonas aeruginosa is essential for multiple virulence factors. Infect Immun 71:1590–5.

28. Bowlin NO, Williams JD, Knoten CA, Torhan MC, Tashjian TF, Li B, Aiello D, Mecsas J, Hauser AR, Peet NP, Bowlin TL, Moir DT. 2014. Mutations in the Pseudomonas aeruginosa needle protein gene pscF confer resistance to phenoxyacetamide inhibitors of the type III secretion system. Antimicrob Agents Chemother 58:2211–20.

29. Rietsch A, Vallet-Gely I, Dove SL, Mekalanos JJ. 2005. ExsE, a secreted regulator of type III secretion genes in Pseudomonas aeruginosa. Proc Natl Acad Sci U S A 102:8006–11.

30. Green ER, Mecsas J. 2016. Bacterial Secretion Systems: An Overview. Microbiol Spectr 4.

31. Avican U, Doruk T, Ostberg Y, Fahlgren A, Forsberg A. 2017. The Tat Substrate SufI Is Critical for the Ability of Yersinia pseudotuberculosis To Cause Systemic Infection. Infect Immun 85.

32. Bageshwar UK, VerPlank L, Baker D, Dong W, Hamsanathan S, Whitaker N, Sacchettini JC, Musser SM. 2016. High Throughput Screen for Escherichia coli Twin Arginine Translocation (Tat) Inhibitors. PLoS One 11:e0149659.

33. Muschiol S, Bailey L, Gylfe A, Sundin C, Hultenby K, Bergstrom S, Elofsson M, Wolf-Watz H, Normark S, Henriques-Normark B. 2006. A small-molecule inhibitor of type III secretion inhibits different stages of the infectious cycle of Chlamydia trachomatis. Proc Natl Acad Sci U S A 103:14566–71.

34. Lee PC, Rietsch A. 2015. Fueling type III secretion. Trends Microbiol 23:296–300.

35. Paul K, Erhardt M, Hirano T, Blair DF, Hughes KT. 2008. Energy source of flagellar type III secretion. Nature 451:489–92.

36. Lee PA, Tullman-Ercek D, Georgiou G. 2006. The bacterial twin-arginine translocation pathway. Annu Rev Microbiol 60:373–95.

37. Troisfontaines P, Cornelis GR. 2005. Type III secretion: More systems than you think. Physiology 20:326–339.

38. Auvray F, Ozin AJ, Claret L, Hughes C. 2002. Intrinsic membrane targeting of the flagellar export ATPase FliI: interaction with acidic phospholipids and FliH. J Mol Biol 318:941–50.

39. Wagner S, Grin I, Malmsheimer S, Singh N, Torres-Vargas CE, Westerhausen S. 2018. Bacterial type III secretion systems: a complex device for the delivery of bacterial effector proteins into eukaryotic host cells. FEMS Microbiol Lett 365.

40. McCaw ML, Lykken GL, Singh PK, Yahr TL. 2002. ExsD is a negative regulator of the Pseudomonas aeruginosa type III secretion regulon. Molecular Microbiology 46:1123–1133.

41. Urbanowski ML, Lykken GL, Yahr TL. 2005. A secreted regulatory protein couples transcription to the secretory activity of the Pseudomonas aeruginosa type III secretion system. Proceedings of the National Academy of Sciences of the United States of America 102:9930–9935.

42. Elwell C, Mirrashidi K, Engel J. 2016. Chlamydia cell biology and pathogenesis. Nat Rev Microbiol 14:385–400.

43. Al-Zeer MA, Al-Younes HM, Kerr M, Abu-Lubad M, Gonzalez E, Brinkmann V, Meyer TF. 2014. Chlamydia trachomatis remodels stable microtubules to coordinate Golgi stack recruitment to the chlamydial inclusion surface. Mol Microbiol 94:1285–97.

44. Derre I. 2015. Chlamydiae interaction with the endoplasmic reticulum: contact, function and consequences. Cell Microbiol 17:959–66.

45. Kumar Y, Valdivia RH. 2008. Actin and intermediate filaments stabilize the Chlamydia trachomatis vacuole by forming dynamic structural scaffolds. Cell Host Microbe 4:159–69.

46. Klockner A, Buhl H, Viollier P, Henrichfreise B. 2018. Deconstructing the Chlamydial Cell Wall. Curr Top Microbiol Immunol 412:1–33.

47. Miller KE. 2006. Diagnosis and treatment of Chlamydia trachomatis infection. Am Fam Physician 73:1411–6.

48. Phillips S, Quigley BL, Timms P. 2019. Seventy Years of Chlamydia Vaccine Research - Limitations of the Past and Directions for the Future. Front Microbiol 10:70.

49. Krupp K, Madhivanan P. 2015. Antibiotic resistance in prevalent bacterial and protozoan sexually transmitted infections. Indian J Sex Transm Dis AIDS 36:3–8.

50. Young BM, Young GM. 2002. YplA is exported by the Ysc, Ysa, and flagellar type III secretion systems of Yersinia enterocolitica. J Bacteriol 184:1324–34.

51. Woehrmann MH, Bray WM, Durbin JK, Nisam SC, Michael AK, Glassey E, Stuart JM, Lokey RS. 2013. Large-scale cytological profiling for functional analysis of bioactive compounds. Mol Biosyst 9:2604–17.

52. Ronald N. Zuckermann JMK, Stephen B. H. Kent and Walter H. Moos. 1992. Efficient method for the preparation of peptoids [oligo(N-substituted glycines)] by submonomer solid-phase synthesis. Journal of the American Chemical Society 114:2.

53. Aiello D, Williams JD, Majgier-Baranowska H, Patel I, Peet NP, Huang J, Lory S, Bowlin TL, Moir DT. 2010. Discovery and characterization of inhibitors of Pseudomonas aeruginosa type III secretion. Antimicrob Agents Chemother 54:1988–99.

54. Kim OK, Garrity-Ryan LK, Bartlett VJ, Grier MC, Verma AK, Medjanis G, Donatelli JE, Macone AB, Tanaka SK, Levy SB, Alekshun MN. 2009. N-Hydroxybenzimidazole Inhibitors of the Transcription Factor LcrF in Yersinia: Novel Antivirulence Agents. Journal of Medicinal Chemistry 52:5626–5634.

55. Enquist PA, Gylfe A, Hagglund U, Lindstrom P, Norberg-Scherman H, Sundin C, Elofsson M. 2012. Derivatives of 8-hydroxyquinoline--antibacterial agents that target intra- and extracellular Gram-negative pathogens. Bioorg Med Chem Lett 22:3550–3.

56. Anantharajah A, Buyck JM, Sundin C, Tulkens PM, Mingeot-Leclercq MP, Van Bambeke F. 2017. Salicylidene Acylhydrazides and Hydroxyquinolines Act as Inhibitors of Type Three Secretion Systems in Pseudomonas aeruginosa by Distinct Mechanisms. Antimicrob Agents Chemother 61.

57. Anantharajah A, Faure E, Buyck JM, Sundin C, Lindmark T, Mecsas J, Yahr TL, Tulkens PM, Mingeot-Leclercq MP, Guery B, Van Bambeke F. 2016. Inhibition of the Injectisome and Flagellar Type III Secretion Systems by INP1855 Impairs Pseudomonas aeruginosa Pathogenicity and Inflammasome Activation. J Infect Dis 214:1105–16.

58. Slepenkin A, Enquist PA, Hagglund U, de la Maza LM, Elofsson M, Peterson EM. 2007. Reversal of the antichlamydial activity of putative type III secretion inhibitors by iron. Infect Immun 75:3478–89.

59. Layton AN, Hudson DL, Thompson A, Hinton JC, Stevens JM, Galyov EE, Stevens MP. 2010. Salicylidene acylhydrazide-mediated inhibition of type III secretion system-1 in Salmonella enterica serovar Typhimurium is associated with iron restriction and can be reversed by free iron. FEMS Microbiol Lett 302:114–22.

60. Hudson DL, Layton AN, Field TR, Bowen AJ, Wolf-Watz H, Elofsson M, Stevens MP, Galyov EE. 2007. Inhibition of type III secretion in Salmonella enterica serovar Typhimurium by small-molecule inhibitors. Antimicrob Agents Chemother 51:2631–5.

61. Morgan JM, Lam HN, Delgado J, Luu J, Mohammadi S, Isberg RR, Wang H, Auerbuch V. 2018. An Experimental Pipeline for Initial Characterization of Bacterial Type III Secretion System Inhibitor Mode of Action Using Enteropathogenic Yersinia. Front Cell Infect Microbiol 8:404.

62. Kauppi AM, Nordfelth R, Uvell H, Wolf-Watz H, Elofsson M. 2003. Targeting bacterial virulence: inhibitors of type III secretion in Yersinia. Chem Biol 10:241–9.

63. Wang D, Zetterstrom CE, Gabrielsen M, Beckham KS, Tree JJ, Macdonald SE, Byron O, Mitchell TJ, Gally DL, Herzyk P, Mahajan A, Uvell H, Burchmore R, Smith BO, Elofsson M, Roe AJ. 2011. Identification of bacterial target proteins for the salicylidene acylhydrazide class of virulence-blocking compounds. J Biol Chem 286:29922–31.

64. Nordfelth R, Kauppi AM, Norberg HA, Wolf-Watz H, Elofsson M. 2005. Small-molecule inhibitors specifically targeting type III secretion. Infect Immun 73:3104–14.

65. Tree JJ, Wang D, McInally C, Mahajan A, Layton A, Houghton I, Elofsson M, Stevens MP, Gally DL, Roe AJ. 2009. Characterization of the effects of salicylidene acylhydrazide compounds on type III secretion in Escherichia coli O157:H7. Infect Immun 77:4209–20.

66. Zambelloni R, Connolly JPR, Huerta Uribe A, Burgess K, Marquez R, Roe AJ. 2017. Novel compounds targeting the enterohemorrhagic Escherichia coli type three secretion system reveal insights into mechanisms of secretion inhibition. Mol Microbiol 105:606–619.

67. Yang F, Korban SS, Pusey PL, Elofsson M, Sundin GW, Zhao Y. 2014. Small-molecule inhibitors suppress the expression of both type III secretion and amylovoran biosynthesis genes in Erwinia amylovora. Mol Plant Pathol 15:44–57.

68. Harmon DE, Davis AJ, Castillo C, Mecsas J. 2010. Identification and characterization of small-molecule inhibitors of Yop translocation in Yersinia pseudotuberculosis. Antimicrob Agents Chemother 54:3241–54.

69. Jessen DL, Bradley DS, Nilles ML. 2014. A type III secretion system inhibitor targets YopD while revealing differential regulation of secretion in calcium-blind mutants of Yersinia pestis. Antimicrob Agents Chemother 58:839–50.

70. Gauthier A, Robertson ML, Lowden M, Ibarra JA, Puente JL, Finlay BB. 2005. Transcriptional inhibitor of virulence factors in enteropathogenic Escherichia coli. Antimicrob Agents Chemother 49:4101–9.

71. Duncan MC, Wong WR, Dupzyk AJ, Bray WM, Linington RG, Auerbuch V. 2014. An NF-kappaB-based high-throughput screen identifies piericidins as inhibitors of the Yersinia pseudotuberculosis type III secretion system. Antimicrob Agents Chemother 58:1118–26.

72. Grishin AV, Luyksaar SI, Kapotina LN, Kirsanov DD, Zayakin ES, Karyagina AS, Zigangirova NA. 2018. Identification of chlamydial T3SS inhibitors through virtual screening against T3SS ATPase. Chem Biol Drug Des 91:717–727.

73. Hao H, Aixia Y, Lei F, Nancai Y, Wen S. 2010. Effects of baicalin on Chlamydia trachomatis infection in vitro. Planta Med 76:76–8.

74. Tsou LK, Lara-Tejero M, RoseFigura J, Zhang ZJ, Wang YC, Yount JS, Lefebre M, Dossa PD, Kato J, Guan F, Lam W, Cheng YC, Galan JE, Hang HC. 2016. Antibacterial Flavonoids from Medicinal Plants Covalently Inactivate Type III Protein Secretion Substrates. J Am Chem Soc 138:2209–18.

75. Guo Z, Li X, Li J, Yang X, Zhou Y, Lu C, Shen Y. 2016. Licoflavonol is an inhibitor of the type three secretion system of Salmonella enterica serovar Typhimurium. Biochem Biophys Res Commun 477:998–1004.

76. Nakasone N, Higa N, Toma C, Ogura Y, Suzuki T, Yamashiro T. 2017. Epigallocatechin gallate inhibits the type III secretion system of Gram-negative enteropathogenic bacteria under model conditions. FEMS Microbiol Lett 364.

77. Zhang Y, Liu Y, Wang T, Deng X, Chu X. 2018. Natural compound sanguinarine chloride targets the type III secretion system of Salmonella enterica Serovar Typhimurium. Biochem Biophys Rep 14:149–154.

78. Choi WS, Lee TH, Son SJ, Kim TG, Kwon BM, Son HU, Kim SU, Lee SH. 2017. Inhibitory effect of obovatol from Magnolia obovata on the Salmonella type III secretion system. J Antibiot (Tokyo) 70:1065–1069.

79. Zhang Y, Liu Y, Qiu J, Luo ZQ, Deng X. 2018. The Herbal Compound Thymol Protects Mice From Lethal Infection by Salmonella Typhimurium. Front Microbiol 9:1022.

80. Felise HB, Nguyen HV, Pfuetzner RA, Barry KC, Jackson SR, Blanc MP, Bronstein PA, Kline T, Miller SI. 2008. An inhibitor of gram-negative bacterial virulence protein secretion. Cell Host Microbe 4:325–36.

81. Bzdzion L, Krezel H, Wrzeszcz K, Grzegorek I, Nowinska K, Chodaczek G, Swietnicki W. 2017. Design of small molecule inhibitors of type III secretion system ATPase EscN from enteropathogenic Escherichia coli. Acta Biochim Pol 64:49–63.

82. Nesterenko LN, Zigangirova NA, Zayakin ES, Luyksaar SI, Kobets NV, Balunets DV, Shabalina LA, Bolshakova TN, Dobrynina OY, Gintsburg AL. 2016. A small-molecule compound belonging to a class of 2,4-disubstituted 1,3,4-thiadiazine-5-ones suppresses Salmonella infection in vivo. J Antibiot (Tokyo) 69:422–7.

83. Zigangirova NA, Kost EA, Didenko LV, Kapotina LN, Zayakin ES, Luyksaar SI, Morgunova EY, Fedina ED, Artyukhova OA, Samorodov AV, Kobets NV. 2016. A small-molecule compound belonging to a class of 2,4-disubstituted 1,3,4-thiadiazine-5-ones inhibits intracellular growth and persistence of Chlamydia trachomatis. J Med Microbiol 65:91–98.

84. Sheremet AB, Zigangirova NA, Zayakin ES, Luyksaar SI, Kapotina LN, Nesterenko LN, Kobets NV, Gintsburg AL. 2018. Small Molecule Inhibitor of Type Three Secretion System Belonging to a Class 2,4-disubstituted-4H-[1,3,4]-thiadiazine-5-ones Improves Survival and Decreases Bacterial Loads in an Airway Pseudomonas aeruginosa Infection in Mice. Biomed Res Int 2018:5810767.

85. Zetterstrom CE, Hasselgren J, Salin O, Davis RA, Quinn RJ, Sundin C, Elofsson M. 2013. The resveratrol tetramer (-)-hopeaphenol inhibits type III secretion in the gram-negative pathogens Yersinia pseudotuberculosis and Pseudomonas aeruginosa. PLoS One 8:e81969.

86. Kang JE, Jeon BJ, Park MY, Yang HJ, Kwon J, Lee DH, Kim BS. 2020. Inhibition of the type III secretion system of Pseudomonas syringae pv. tomato DC3000 by resveratrol oligomers identified in Vitis vinifera L. Pest Manag Sci 76:2294–2303.

87. Lv Q, Li S, Wei H, Wen Z, Wang Y, Tang T, Wang J, Xia L, Deng X. 2020. Identification of the natural product paeonol derived from peony bark as an inhibitor of the Salmonella enterica serovar Typhimurium type III secretion system. Appl Microbiol Biotechnol 104:1673–1682.

88. Lv Q, Chu X, Yao X, Ma K, Zhang Y, Deng X. 2019. Inhibition of the type III secretion system by syringaldehyde protects mice from Salmonella enterica serovar Typhimurium. J Cell Mol Med 23:4679–4688.

89. Li J, Sun W, Guo Z, Lu C, Shen Y. 2014. Fusaric acid modulates Type Three Secretion System of Salmonella enterica serovar Typhimurium. Biochem Biophys Res Commun 449:455–9.

90. Li J, Lv C, Sun W, Li Z, Han X, Li Y, Shen Y. 2013. Cytosporone B, an inhibitor of the type III secretion system of Salmonella enterica serovar Typhimurium. Antimicrob Agents Chemother 57:2191–8.

91. Kimura K, Iwatsuki M, Nagai T, Matsumoto A, Takahashi Y, Shiomi K, Omura S, Abe A. 2011. A small-molecule inhibitor of the bacterial type III secretion system protects against in vivo infection with Citrobacter rodentium. J Antibiot (Tokyo) 64:197–203.

92. Ma YN, Chen L, Si NG, Jiang WJ, Zhou ZG, Liu JL, Zhang LQ. 2019. Identification of Benzyloxy Carbonimidoyl Dicyanide Derivatives as Novel Type III Secretion System Inhibitors via High-Throughput Screening. Front Plant Sci 10:1059.

93. Wagener BM, Anjum N, Evans C, Brandon A, Honavar J, Creighton J, Traber MG, Stuart RL, Stevens T, Pittet JF. 2020. Alpha-tocopherol Attenuates the Severity of Pseudomonas aeruginosa-induced Pneumonia. Am J Respir Cell Mol Biol doi:10.1165/rcmb.2019-0185OC.

94. Liu Y, Zhang Y, Zhou Y, Wang T, Deng X, Chu X, Zhou T. 2019. Cinnamaldehyde inhibits type three secretion system in Salmonella enterica serovar Typhimurium by affecting the expression of key effector proteins. Vet Microbiol 239:108463.

95. Bliska JB, Guan KL, Dixon JE, Falkow S. 1991. Tyrosine Phosphate Hydrolysis of Host Proteins by an Essential Yersinia-Virulence Determinant. Proceedings of the National Academy of Sciences of the United States of America 88:1187–1191.

96. Rabin SD, Hauser AR. 2005. Functional regions of the Pseudomonas aeruginosa cytotoxin ExoU. Infect Immun 73:573–82.

97. Rangel SM, Diaz MH, Knoten CA, Zhang A, Hauser AR. 2015. The Role of ExoS in Dissemination of Pseudomonas aeruginosa during Pneumonia. PLoS Pathog 11:e1004945.

98. Portnoy DA, Moseley SL, Falkow S. 1981. Characterization of plasmids and plasmid-associated determinants of Yersinia enterocolitica pathogenesis. Infect Immun 31:775–82.

99. Winter SE, Winter MG, Poon V, Keestra AM, Sterzenbach T, Faber F, Costa LF, Cassou F, Costa EA, Alves GE, Paixao TA, Santos RL, Baumler AJ. 2014. Salmonella enterica Serovar Typhi conceals the invasion-associated type three secretion system from the innate immune system by gene regulation. PLoS Pathog 10:e1004207.

100. Matsui H, Eguchi M, Ohsumi K, Nakamura A, Isshiki Y, Sekiya K, Kikuchi Y, Nagamitsu T, Masuma R, Sunazuka T, Omura S. 2005. Azithromycin inhibits the formation of flagellar filaments without suppressing flagellin synthesis in Salmonella enterica serovar typhimurium. Antimicrob Agents Chemother 49:3396–403.

